# Game structure shapes normative preferences and inter-brain neural synchrony in free-allocation public goods games

**DOI:** 10.64898/2026.11.10.765617

**Authors:** Yang Li, Weijie Xing, Yifan Wu, Xin Wang, Wucheng Dong, Xingru Chen, Longzhao Liu, Hongwei Zheng, Shaoting Tang

## Abstract

Threshold public goods games (TPGGs) are generally expected to facilitate coordination more effectively than linear public goods games (LPGGs), because an explicit provision point can support non-zero equilibria and transform a social dilemma into a coordination problem. This prediction, however, is based largely on settings in which group members contribute to a single shared pool. Whether it extends to environments in which individuals freely allocate resources across multiple partners remains unknown. Here, using a three-person ring network and functional near-infrared spectroscopy hyperscanning, we examined how free allocation shaped behavioral and neural signatures in the two game structures. Contrary to conventional expectations, the LPGG produced stronger partner-specific reciprocity and greater alignment to history-dependent contribution standards than the TPGG. This behavioral differentiation was accompanied by stronger dyadic inter-brain synchrony in the dorsomedial prefrontal cortex during the LPGG. The two structures also elicited distinct patterns of individual neural activity and dmPFC–rTPJ functional connectivity. These findings suggest that the expected coordination advantage of threshold public goods games may not generalize to settings that allow partner-specific resource allocation, highlighting the importance of allocation flexibility in shaping the relationship between game structure and cooperative behavior.

## Introduction

Shared consensus can sustain cooperation over repeated interactions [5–7]. Many real-world collective-action problems also involve explicit provision targets, such that collective benefits are realized only when aggregate contributions reach a predetermined threshold. In threshold public goods games (TPGGs), this provision point provides a shared coordination reference around which group members can form expectations, adjust their contributions and, through repeated interaction, converge on a shared contribution standard [8–10]. Linear public goods games (LPGGs), by contrast, generate continuous returns from every unit contributed but, when the marginal per-capita return is below one, retain zero contribution as the unique one-shot Nash equilibrium. From the perspective of interdependence theory [2, 4, 11], the LPGG therefore exposes a direct conflict between individual and collective incentives, whereas the TPGG can transform this social dilemma into a coordination problem centred on a common target. This distinction motivates the prevailing theoretical expectation that TPGGs should promote behavioral convergence, collective coordination and norm alignment more effectively than LPGGs [1–4, 12].

However, this classical account rests on a restrictive assumption: all group members contribute to a single shared public pool, and their contributions are aggregated into a common outcome [13]. Many real-world collective behavior are organized differently. In organizations, teams, scientific collaborations [14], community governance and digital platforms, individuals often interact with several partners simultaneously [15]. Cooperation therefore requires decisions not only about how much to contribute, but also about how limited effort and resources should be allocated across partners. A team member may, for example, concentrate effort on a project involving a particular collaborator or divide resources among several collaborative ties [16]. Collective outcomes in such settings emerge from multiple interconnected interactions rather than from a single pool. Despite its real-world relevance, it is unclear whether allowing partner-specific allocation preserves or reverses the behavioral differences conventionally expected between LPGGs and TPGGs.

Hyperscanning offers a means of examining these interactive processes directly [17]. Reciprocal cooperation is not simply the aggregate of independent decisions; it unfolds through repeated exchanges in which individuals monitor others’ behavior, respond to feedback and adjust their subsequent actions [18–20]. By recording neural responses simultaneously from interacting individuals, hyperscanning enables cooperation to be studied in terms of interpersonal neural coupling alongside individual neural signatures [21, 22]. Reviews and meta-analyses indicate that cooperative interaction is frequently accompanied by increased inter-brain neural synchrony (INS) in cortical regions implicated in social cognition, potentially reflecting shared attention, mutual prediction and interpersonal coordination [23–26]. Hyperscanning is therefore particularly well suited to testing whether free allocation engages distinct interactive processes in LPGGs and TPGGs [27, 28].

If free allocation affects only the payoff structure of LPGGs and TPGGs, without altering how individuals evaluate and respond to their partners, its consequences may be limited to behavioral outcomes. However, if free allocation also influences partner monitoring, norm alignment, and reciprocal adjustment, the two game structures may elicit distinct neural signatures [29, 30], particularly in dyadic INS. The dorsomedial prefrontal cortex (dmPFC) provides a plausible neural substrate for these processes because of its involvement in social inference [31], value computation [32], and norm processing [33, 34]. Accordingly, enhanced INS in prefrontal regions may reflect how individuals predict their partners’ behavior and adjust their decisions during social interaction [35].

We investigated these questions using free-allocation public goods games implemented in a minimal three-person ring network. Each participant belonged simultaneously to two dyadic public-goods pools and allocated a limited endowment between two partners [36]. The resulting task was not a collection of isolated dyadic games: every participant was embedded in two concurrent relationships, and decisions directed towards one partner constrained the resources available for the other [37]. This structure allowed collective cooperation to be decomposed into partner-specific allocation processes. Within the same network architecture, we compared the LPGG and TPGG and combined behavioral modelling with functional near-infrared spectroscopy (fNIRS) hyperscanning. This approach enabled us to test whether game structure shaped contribution dynamics and norm alignment, as well as dyadic INS, individual neural activity and intra-brain functional connectivity.

At the behavioral level, we examined how partner-specific contribution differences evolved across rounds and estimated conditional contribution patterns from individuals’ own and their partners’ preceding behavior [38]. The corresponding coefficients captured behavioral stickiness and imitation. We then fitted norm–utility models to estimate individual alignment to history-dependent contribution standards, permitting a quantitative comparison of normative preference weight between game structures [39, 40]. At the neural level, we measured changes in oxygenated haemoglobin concentration during live interaction. Dyadic INS within each public-goods pool was quantified using wavelet transform coherence. We asked whether the LPGG and TPGG elicited different levels of dmPFC synchrony and whether this synchrony was associated with model-derived normative preference weight or with partner-specific contribution differences. We additionally analyzed individual neural activity and dmPFC–rTPJ functional connectivity to determine whether game structure differentiated neural responses at the individual level.

Our findings argue that explicit thresholds may not necessarily strengthen coordination and norm alignment in public goods games. The coordination effects of a threshold depend on whether contributions enter a single shared pool or can be allocated across multiple partner-specific pools. Under free allocation, the TPGG threshold no longer provides a single collective standard and may instead create multiple local coordination risks [41]. By contrast, the continuous feedback generated by the LPGG preserves information about each partner’s behavior, supporting partner-specific reciprocity and stronger alignment to history-dependent contribution standards. Consistent with this account, the LPGG elicited stronger dyadic dmPFC INS; critically, the stronger synchrony was associated with contribution differences rather than stronger alignment to established norms. Dyadic INS therefore appeared to reflect the ongoing coordination demands of reciprocal monitoring and norm updating, rather than a stable state of behavioral consensus [42]. By integrating behavioral modelling and hyperscanning, our study shows how freedom over resource allocation can reorganize the behavioral and neural mechanisms of cooperation across interconnected social relationships.

## Methods

### Participants

We recruited 132 undergraduate students from Beihang University. Two groups (six participants) were excluded following participant withdrawal during the experiment, leaving 126 participants in 42 three-person groups for the behavioral analyses (75 female; age range, 17–25 years). Participants received performance-based cash payments of US$30–35. Two additional groups were excluded from the neural analyses because of fNIRS recording problems, resulting in a neural sample of 120 participants in 40 groups.

All participants were right-handed, had normal or corrected-to-normal vision and possessed basic arithmetic skills. They completed comprehension control questions to assess their understanding of the task. The study was approved by the Beihang University Biomedical Ethics Committee (protocol no. BM20250183) and conducted in accordance with the Declaration of Helsinki. Written informed consent was obtained from all participants before the experiment.

### Procedure

Participants completed a repeated contribution task involving three dyadic public pools in either the LPGG or TPGG condition. Each session involved three previously unacquainted participants who interacted anonymously and remained in the same group throughout the experiment. Participants’ identities were not disclosed to other group members.

At the beginning of each session, each participant was randomly assigned a unique identification code that remained fixed throughout the task and did not reveal their real identities. To limit communication and social cues outside the task, participants were instructed to remain silent, attend only to their own computer screens and refrain from observing other participants or their displays. To minimize motion artefacts during concurrent fNIRS recordings, participants were instructed to operate the mouse with their right hand and keep their left hand resting on the table. Keyboard use was prohibited. Mouse movements and clicks were permitted only during the operation phases, and participants were instructed to remain still during all other periods.

### Pre-experiment instructions and comprehension check

The day before the main experiment, registered participants attended a preliminary session to complete a pre-experiment questionnaire assessing eligibility and collecting background information for subsequent analyses (Supplementary Section. 5). The questionnaire included demographic information, measures of incentive economic preferences and social value orientation (SVO). Demographic information comprised age, gender and academic discipline. Economic preferences were elicited using decision tasks assessing risk aversion, patience, trust, altruism, and positive and negative reciprocity [67]. SVO was assessed using the SVO slider task, in which participants selected allocations of resources between themselves and another person [68]. Eligible participants read the experimental instructions and provided written informed consent. They were also informed that they could participate in the experiment only once.

Before the main task, participants received detailed on-screen instructions describing the task structure, contribution rules, payoff calculations and timing of each round for their assigned game condition (LPGG or TPGG). Participants then completed control questions requiring them to calculate payoffs for specified contribution profiles (Supplementary Section. 7). The task began only after all participants had answered every question correctly. This procedure was intended to minimize misunderstanding of the task rules as a source of confusion in contribution decisions [69].

### Free-allocation task

At the beginning of each round, participants received an initial endowment of 35 wealth units. Using an on-screen slider with eight discrete settings, they allocated this endowment between two dyadic public pools, each shared with one of the other two group members. Any unallocated wealth was retained. Total contributions across the two pools could not exceed 35 units.

In the LPGG, the combined contributions to each pool were multiplied by 1.5 and divided equally between its two members. Each unit contributed therefore generated 1.5 units in collective returns, of which the contributor received 0.75 units. This payoff structure created a social dilemma: contributions increased collective payoffs but reduced the contributor’s own payoff, holding the partner’s contribution constant. In the TPGG, the combined contributions to each pool were multiplied by 1.5 and compared with a threshold. If the resulting amount met or exceeded the threshold, each member received half the threshold value. Otherwise, neither member received a return from that pool, and their contributions were forfeited. Returns were thus conditional on threshold attainment and capped at the threshold value. Because contributions that failed to meet the threshold yielded zero returns, this mechanism introduced a non-linear risk–reward structure. Three nominal thresholds were used in the experiment: 30, 45 and 60 units, corresponding to contributions of 10, 15 and 20 units per person, respectively, when both partners contributed equally. Threshold levels were presented in a randomised, balanced sequence, with each level appearing twice during the adaptation stage and five times during the task stage. On each presentation, the nominal threshold was decreased by 4 units with probability 0.25, left unchanged with probability 0.50, or increased by 4 units with probability 0.25.

Each round followed a fixed sequence. A fixation cross was presented for 5 s, followed by a 12-s operation phase during which participants selected their contributions to dyadic communities using the mouse. A 3-s interval allowed the system to calculate and record outcomes, after which feedback was displayed for 10 s. In the LPGG, feedback showed the combined contributions to each pool, the total return after multiplication and the amount distributed to each member. In the TPGG, feedback additionally indicated whether each pool had reached its threshold and the corresponding return. A subsequent 5-s screen displayed each participant’s final wealth for that round, comprising their retained endowment and returns from both pools. Successive rounds were separated by a 10-s rest period, during which participants were instructed to remain still. The experimental session lasted approximately 30 min.

### Post-experiment measurement

After the experiment, participants completed a questionnaire assessing their expectations about contributions and the motives underlying their contribution decisions, alongside feedback on the task (Supplementary Section. 6).

### Behavioral data acquisition and model-free analysis

We developed a custom web-based platform to administer the task, record interactions in real time and align behavioral events with fNIRS recordings. The platform supported simultaneous, anonymous participation by three individuals. The front end was built using Vue.js, and trial-by-trial behavioral data, including individual contribution decisions, were stored in a MySQL database. WebSocket communication coordinated task events between the server and participant clients, maintaining a common trial sequence across the group. Each task transition was logged with a millisecond-resolution timestamp. These timestamps were subsequently used to align behavioral events with the fNIRS recordings and define event markers for neural analyses.

### Round-level analysis

We conducted round-level analyses to characterize contribution dynamics under free allocation in the LPGG and TPGG. For each participant and round, contribution differentiation was defined as the absolute difference between their contributions to the two dyadic public pools:

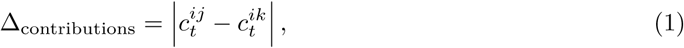

where *c_t_^ij^* and *c_t_^ik^* denote participant *i*’s contributions to the pools shared with partners *j* and *k* as in main text, respectively, in round *t*. Higher values indicate greater differentiation in allocation between partners.

For each group and round, payoff inequality was defined as the mean absolute payoff difference across the three unique pairs of group members:

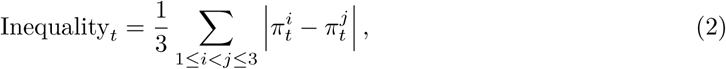

where *π_t_^i^* denotes the final payoff of individual *i* in round *t*. Higher values indicate greater within-group payoff inequality.

To assess temporal trends, we fitted linear mixed-effects models with round as a fixed effect. Models of individual-level outcomes, including total contributions and contribution differentiation, included a random intercept for participant identity, whereas models of within-group payoff inequality included a random intercept for group identity. Models were fitted separately for the LPGG and TPGG:

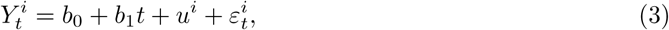

where *Y_t_^i^*denotes the behavioral outcome for participant *i* in round *t*, *u^i^*is the participant-specific random intercept and *ε_t_^i^* is the residual error.

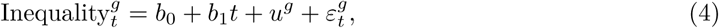

where Inequality*_t_^g^*denotes payoff inequality within group *g* in round *t*, *u^g^*is the group-specific random intercept and *ε_t_^g^* is the residual error.

We further compared the LPGG and TPGG in each round to determine when behavioral differences emerged under free allocation. Between-condition comparisons were conducted for contribution differentiation. The resulting *P* values were adjusted across rounds using the FDR procedure.

### Individual-level analysis

To characterize how free allocation shaped partner-specific contribution strategies, we analyzed contributions at the level of dyadic public pools. In each three-person group, participants formed three dyadic pools, (1, 2), (1, 3) and (2, 3). In each round, a focal participant *i* could freely allocate their endowment across the two dyadic pools to which they belonged. Thus, each observation was defined by a focal individual, a partner *j*, a dyadic public pool (*i, j*) and a round *t*.

We fitted Bayesian linear mixed-effects models to estimate conditional contribution patterns. The dependent variable was *c_t_^ij^*. The predictors included the focal participant’s contribution to the same pool in the preceding round, *c_t-1_^ij^*, and the partner’s preceding contribution to that pool, *c_t-1_^ji^*. Their respective coefficients were termed *stickiness* and *imitation*, capturing the extent to which current contributions were associated with the focal participant’s own and their partner’s previous contributions. We also included the interaction between the two lagged contributions. The models contained participant-specific random intercepts and random slopes for both lagged contribution terms and were fitted separately for the LPGG and TPGG:

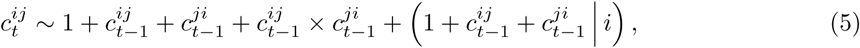

where the fixed-effect coefficients of *c_t-1_^ij^* and *c_t-1_^ji^* quantify stickiness and imitation, respectively.

As a robustness analysis, we fitted an alternative specification in which the random effects were defined at the dyadic-pool level:

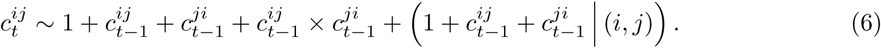

This specification tested whether the estimated patterns of stickiness and imitation were robust to modelling heterogeneity between dyadic public pools rather than between individual participants.

### Norm-utility model

To quantify the extent to which allocation decisions were constrained by history-dependent contribution standards, we fitted separate norm–utility models for the LPGG and TPGG. Individual *i* interacted with two partners, *j* and *k*, received an endowment of *e* = 35 tokens per round and selected an allocation **c***_t_^i^* = (*c_t_^ij^_t_, c_t_^ik^*) from the feasible decision set

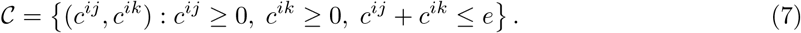

Given the discrete allocation increments used in the experiment, C contained 36 feasible options. For each dyadic public pool, we defined a history-dependent contribution standard. In the LPGG, the standard for pool (*i, j*) in round *t*, denoted *c_t_*^∗^*^,ij^*, was the mean contribution of its two members over the preceding *h* rounds:

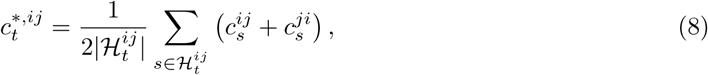

where

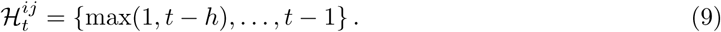

Thus, *c_t_*^∗*,ij*^ represented the recent contribution standard established jointly within a specific dyadic pool, rather than the focal participant’s own contribution history alone.

For the TPGG, contribution standards were conditioned on threshold level. Let *ℓ*(threshold*_t_*) denote the level of the threshold threshold*_t_* assigned to pool (*i, j*) in round *t*. The relevant history comprised the *h* most recent preceding rounds in which the same dyadic pool faced the same threshold level:

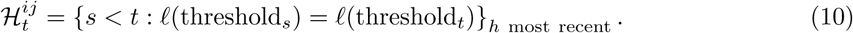

We set *h* = 3, thereby capturing recent contribution standards while limiting the influence of more distant interactions that might no longer guide current allocation decisions.

For each candidate allocation (*c^ij^, c^ik^*) ∈ C, we first calculated the corresponding material payoff. The squared deviation of the candidate allocation from the history-dependent contribution standards was defined as

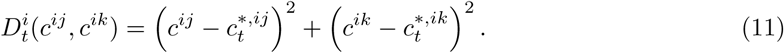

The utility associated with each candidate allocation was then specified as

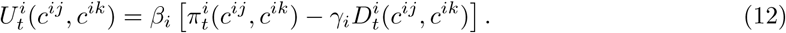

Here, *γ_i_* quantifies individual alignment to history-dependent contribution standards. Positive values indicate that participant *i* assigns lower utility to allocations that deviate from these standards, whereas values close to zero indicate weak norm alignment. The inverse-temperature parameter *β_i_ >* 0 determines the extent to which decisions are concentrated around higher-utility allocations.

Observed decisions were linked to utility using a softmax rule. The probability that individual *i* selected the observed allocation **c***_t_^i,^*^obs^ = (*c_t_^ij^, c_t_^ik^*) in round *t* was defined as

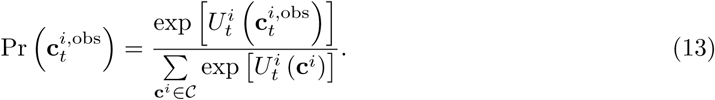

The likelihood of the complete sequence of observed allocations was therefore

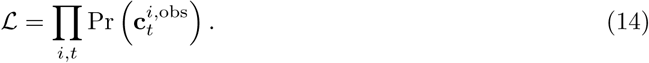

Parameters were estimated in a Bayesian framework, with separate models fitted to each three-person group. Each participant was assigned a normative preference weight, *γ_i_*, and a inverse-temperature parameter, *β_i_*. We specified the priors

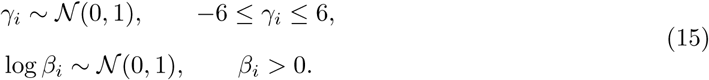

Posterior distributions were estimated using Markov chain Monte Carlo sampling with four chains, each comprising 1,000 warm-up iterations and 2,000 post-warm-up iterations. Convergence and sampling efficiency were assessed using *R̂* and the bulk and tail effective sample sizes.

To compare parameter estimates between LPGG and TPGG, we independently resampled the individual-level posterior means with replacement within the LPGG and TPGG and calculated the difference between the resulting mean estimates:

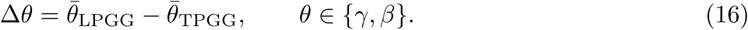

The resulting bootstrap distributions quantified uncertainty in the between-structure differences based on individual-level posterior point estimates.

### fNIRS data acquisition

Neural data from all three participants were acquired simultaneously using the ETG-4100 optical topography system (Hitachi Medical Corporation), sampling at 10 Hz and employing near-infrared wavelengths of 695 nm and 830 nm. Three identical customized optode probe sets were used, each comprising two sets of probes. Optode positions were localised according to the international 10-10 system using the fNIRS Optodes’ Location Decider [70]. The first probe set, comprising five emitters and four detectors, covered the dorsal medial prefrontal cortex, with its optode centred at Fpz. The second probe set was positioned over the right temporoparietal junction, aligned so that the emitter and detector were placed at CP6 and TP8 (Supplementary Fig. 6). Before every session, all optode arrays were inspected and repositioned as needed to ensure consistent placement both within and across participant groups. Following the modified Beer-Lambert law, concentration changes in HbO and HbR were extracted from each channel. Because HbO is the most sensitive haemodynamic variable and shows a strong positive correlation with the BOLD signal, only HbO concentration changes were analyzed.

### fNIRS data processing

#### Data preprocessing

We characterized neural responses at three levels: (1) inter-brain neural synchrony (INS), quantified as dyadic neural coupling; (2) individual neural activity; and (3) intra-brain functional connectivity between the dmPFC and rTPJ. Raw fNIRS signals were detrended using a discrete cosine transform high-pass filter with a 128-s cut-off period. Temporal autocorrelation was subsequently corrected using a pre-colouring procedure with a filter derived from the HRF transfer function [71].

Analysis windows for individual neural responses were defined according to the temporal structure of the experimental task. We defined the selection period (Supplementary Fig. 7a) as the operation phase during which participants evaluated the available options and submitted their decisions. This interval served as the primary task epoch for the analysis of decision-related neural responses.

#### Inter-brain neural synchrony

We used wavelet transform coherence (WTC) to quantify neural coupling between pairs of preprocessed fNIRS time series. To identify the frequency of interest (FOI), we first averaged WTC across all participant pairs, 127 frequency bins and 13 dmPFC and rTPJ channels, without distinguishing between game structures. Task-related coherence was calculated by subtracting mean resting-state coherence from mean coherence during the operation phase. Relative to rest, coherence increased significantly within the frequency range of 0.0288–0.0384 Hz, corresponding to periods of approximately 25–35 s. This range was consistent with the temporal structure of the behavioral task (Supplementary Fig. 7b) and was lower than frequencies typically dominated by head motion, respiration (∼ 0.2–0.3 Hz) and cardiac activity (∼ 1 Hz). Subsequent INS analyses were therefore restricted to this frequency band.

For each channel, Fisher-transformed coherence values within the FOI were averaged over the operation phase and corrected for resting-state coherence to obtain a mean task-related INS value for each dyad. Two-sided one-sample *t* tests were used to determine whether task-related INS differed from zero within each game structure. To identify structure-dependent differences, mean INS was further compared between the LPGG and TPGG using channel-wise two-sample *t* tests.

To determine whether the observed effects were specific to genuine social interaction, we performed a non-parametric pseudo-group permutation analysis separately within each game structure. In each of 1,000 permutations, participants were reassigned to pseudo-groups while preserving their original game structure. Participants from the same real group were constrained to different pseudo-groups, ensuring that no pseudo-group contained more than one member of an original interacting group (Fig. 3d). INS was then recalculated using the same procedure applied to the real groups. For each effect of interest, the test statistic obtained from the real groups, *T*_0_, was compared with the corresponding null distribution, *T*_1_*, T*_2_*, . . ., T_B_*, generated from the pseudo-groups. For a two-sided test, the permutation *P* value was calculated as

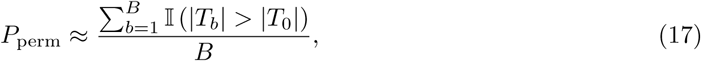

where I denotes the indicator function and *B* = 1,000. This procedure was applied both to task-related INS within each structure and to the LPGG–TPGG difference.

To examine the relationship between INS and behavioral dynamics, we calculated two dyad-level measures. *γ* for dyad (*i, j*) was defined as the mean of the two partners’ individual estimates:

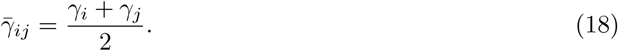

Contribution difference was defined as the mean absolute difference between the partners’ contributions to their shared dyadic pool across rounds:

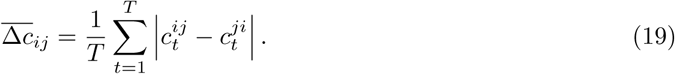

These measures were entered separately into regression models predicting averaged dyadic INS for each channel.

#### Neural activity

The initial 15-s waiting period preceding each operation phase served as the baseline (Fig. 5a). Task-related neural activity was quantified as the change in oxygenated haemoglobin (oxy-Hb) concentration during the operation phase relative to this baseline. For each trial and channel, oxy-Hb signals during the operation phase were standardized using the mean and standard deviation of the corresponding baseline period.

We first averaged task-related neural activity across rounds for each participant and compared participant-level mean activity between the LPGG and TPGG. To assess whether neural activity tracked collective behavioral dynamics, we then averaged neural activity and total contributions across participants within each round and regressed round-level mean neural activity against the corresponding mean contribution. Finally, we examined within-participant covariation between neural activity and behavior by correlating each participant’s round-by-round total contributions with their corresponding round-by-round neural activity separately for each channel. The resulting correlation coefficients were Fisher *z*-transformed and tested against zero using two-sided one-sample *t* tests.

#### Intra-brain functional connectivity

Intra-brain functional connectivity between the dmPFC and rTPJ was quantified using wavelet transform coherence (WTC). For each participant, coherence was calculated between each of the 12 dmPFC channels and the rTPJ channel, yielding 12 dmPFC–rTPJ channel pairs. For each round and channel pair, coherence values were averaged over the operation phase and Fisher *z*-transformed to provide an index of task-related functional connectivity.

Following the neural activity analysis, we examined whether dmPFC–rTPJ connectivity tracked within-participant behavioral dynamics. For each participant and channel pair, we correlated round-by-round total contributions with the corresponding round-by-round functional-connectivity values. The resulting correlation coefficients were Fisher *z*-transformed and tested against zero using two-sided one-sample *t* tests.

#### Analysis plan

All statistical analyses and final visualizations were performed using R version 4.3.1 and MATLAB R2022a.

Bayesian linear mixed-effects models were fitted using the brms package with its default priors [72]. Continuous predictors were standardized to a mean of 0 and a standard deviation of 1 before model fitting. For each model, we report the posterior means of the regression coefficients and correlation estimates, together with highest-density intervals calculated using the bayestestR package (version 0.15.2). Posterior distributions were estimated using Markov chain Monte Carlo sampling with 10 chains, each comprising 1,000 warm-up iterations and 3,000 post-warm-up iterations. Convergence and sampling efficiency were evaluated using the potential scale reduction statistic, *R̂*, and the bulk and tail effective sample sizes (Bulk-ESS and Tail-ESS). Across all models, *R̂* did not exceed 1.05, and the effective sample sizes were sufficient to support stable posterior estimation.

The Bayesian norm–utility model used to estimate the normative preference weight was implemented in Stan and fitted using the cmdstanr package. Models were fitted separately to each three-person group, with individual-specific parameters estimated for the normative preference weight, *γ_i_*, and inverse temperature, *β_i_*. We specified the priors

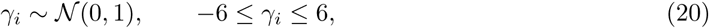

and

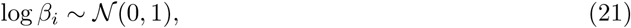

with the latter constraining *β_i_*to positive values. Posterior distributions were estimated using four Markov chains, each comprising 1,000 warm-up iterations and 2,000 post-warm-up iterations. Posterior summaries for *γ_i_* and *β_i_* included the mean, median, standard deviation, median absolute deviation, 5th and 95th percentiles, *R̂*, Bulk-ESS and Tail-ESS. The posterior mean of *γ_i_* was subsequently used as an individual-level index of normative preference weight in the behavioral and neural analyses.

Raw fNIRS signals were preprocessed in MATLAB using functions based on the NIRS-SPM package [71]. Wavelet coherence between pairs of neural time series was calculated using the WTC implementation in MATLAB. Statistical comparisons of neural measures within and between the LPGG and TPGG were performed separately for each channel or channel pair, as appropriate. Multiple comparisons across channels were controlled using false-discovery-rate correction.

## Results

### Linear and threshold public goods games with free-allocation mechanism

We considered a minimal three-person network environment, the smallest setting in which individuals can allocate resources across more than one public good (Fig.1a). In each group, three dyadic public pools are formed, one for each unordered pair of individuals: (1, 2), (1, 3) and (2, 3). Unlike the previous work about PGGs on networks, individuals are not assigned to a fixed public pool. Instead, each individual freely allocated a finite endowment across the two dyadic pools to which they belong, thereby selecting both the amount contributed and the social target of the contribution. Thus, the payoff of an individual is the sum of two components: the reservation in the private account (Fig.1b) and the return from the two dyadic pools. Formally, we set individual 1 as the focal individual. Let *e* denote the endowment of each individual. Individual 1 contributes *c*_12_ to the pool shared with individual 2, and *c*_13_ to the pool shared with individual 3. The feasible allocation set is therefore

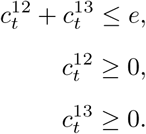

In the linear public goods game (LPGG), contributions to each dyadic pool are multiplied by a productivity factor *P* and then shared equally between the individuals in that pool (Fig.1b, left panel). So the final payoff of individual 1 is

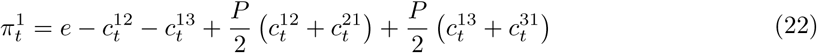

Here, *c*_21_ is individual 2’s contribution to the pool (1, 2); *c*_31_ is individual 3’s contribution to the pool (1, 3).

In the threshold public goods game (TPGG), the payoff-generating rule is no longer proportional to the total contributions. For each pool, the total contributions are first multiplied by the productivity factor *P* . The resulting public output is then compared with a pre-specified *threshold*. If the public output reaches or exceeds it, the pool generates a fixed return equal to *threshold*, which is shared equally between the individuals in that pool. Otherwise, the pool generates no public return (Fig.1b, right panel). Contributions above the threshold do not increase the public return. The payoff of individual 1 is therefore

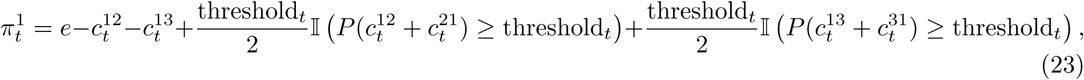

where I(·) is an indicator function that equals 1 when the corresponding dyadic pool reaches the threshold and 0 otherwise.

The experiment comprised 21 rounds: a 6-round adaptation stage followed by a 15-round main task stage. Participants were assigned to one fixed game structure, either the LPGG or the TPGG. The productivity factor *P* was fixed at 1.5. In each round, participants used two sliders to allocate an initial endowment of 35 points, with contributions constrained to increments of 5 points (Fig.1a, c). Throughout the experiment, cerebral haemodynamic activity in predefined regions of interest was continuously recorded using multi-channel fNIRS.

### Free allocation mechanism shapes different behavioral dynamics between LPGG and TPGG

We first examined how individuals differentiated their contributions between the two dyadic public pools over rounds. We fitted linear mixed-effects models to the absolute difference between each individual’s contributions to the two pools, with round as a fixed effect and individual identity as a random intercept (Fig. 1d, f). Contribution differences increased over rounds in the LPGG (*β* = 0.390, SE = 0.029, *t*(1259) = 13.24, *P <* 2.2 × 10^−16^), whereas there was no evidence of a temporal trend in the TPGG (*β* = −0.015, SE = 0.033, *t*(1259) = −0.45, *P* = 0.656). Round-wise comparisons further showed that contribution differences were significantly greater in the LPGG than in the TPGG at multiple rounds beyond the initial rounds (Fig. 1f, red error bars; two-sample *t* tests with FDR correction). Together, these findings indicate increasing differentiation of contributions between partners in the LPGG, consistent with a greater role for partner-specific allocation in this game structure.

**Fig. 1:**
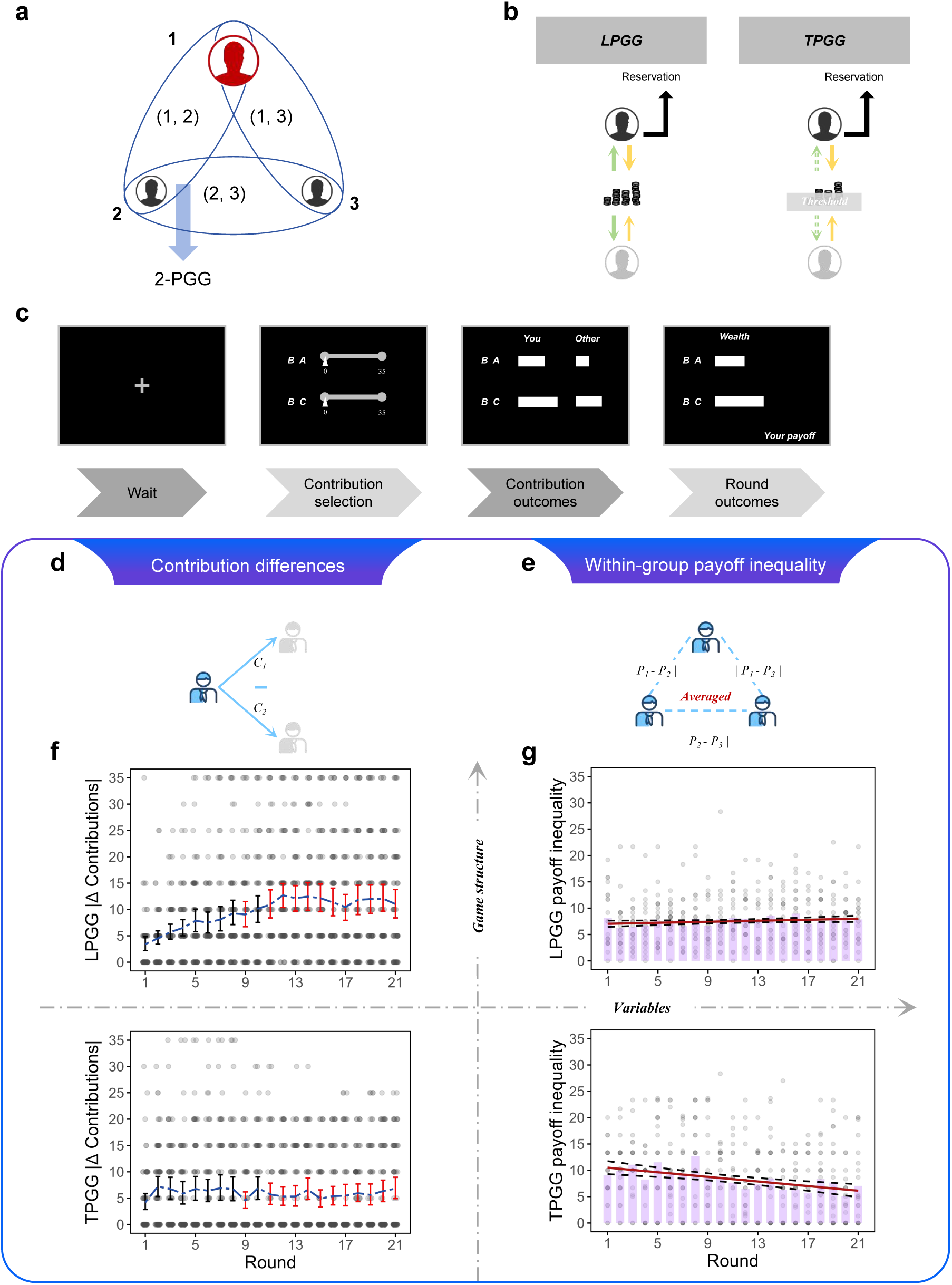
Task structure and behavioral dynamics under free allocation. **a,** Three-person ring network comprising three dyadic public pools. **b,** Payoff structures of the LPGG and TPGG. **c,** Trial sequence, including contribution selection, contribution feedback and the presentation of round-level outcomes. **d,e,** Definitions of partner-specific contribution differences (**d**) and within-group payoff inequality (**e**). **f,g,** Changes in contribution differences (**f**) and payoff inequality (**g**) across rounds. Under the LPGG, contribution differences between partners increased progressively and payoff inequality tended to increase, whereas the TPGG showed no comparable increase in partner-specific allocation and a progressive reduction in payoff inequality.

We next examined how within-group inequality in final payoffs changed over rounds (Fig. 1e, g). In the LPGG, payoff inequality showed a positive but non-significant temporal trend (*β* = 0.047, SE = 0.024, *t*(19) = 1.96, *P* = 0.0653). By contrast, payoff inequality decreased over rounds in the TPGG (*β* = −0.219, SE = 0.050, *t*(19) = −4.36, *P* = 3.37 × 10^−4^). These patterns suggest that free allocation in the LPGG may have facilitated more selective reciprocity towards specific partners, potentially contributing to the observed upward trend in payoff inequality. By contrast, individuals in the TPGG exhibited less differentiation in their contributions between partners, with payoff inequality decreased over rounds.

### The LPGG elicited stronger norm-based coordination under free allocation

We next examined conditional contribution patterns in the dyadic public pools to characterize how individuals adjusted their decisions in response to their own and their partners’ previous contributions (Fig. 2a). We modelled the focal individual’s contribution to a given pool as a function of their own contribution and their partner’s contribution to the same pool in the preceding round [43–45]. We termed the corresponding regression coefficients *stickiness* and *imitation*, respectively, reflecting stickiness to individuals’ own contributions and influence by their partners’ contributions [38]. We also included an interaction between the two lagged contribution terms and fitted a Bayesian recursive model with individual-specific random intercepts and random slopes for both lagged predictors.

**Fig. 2:**
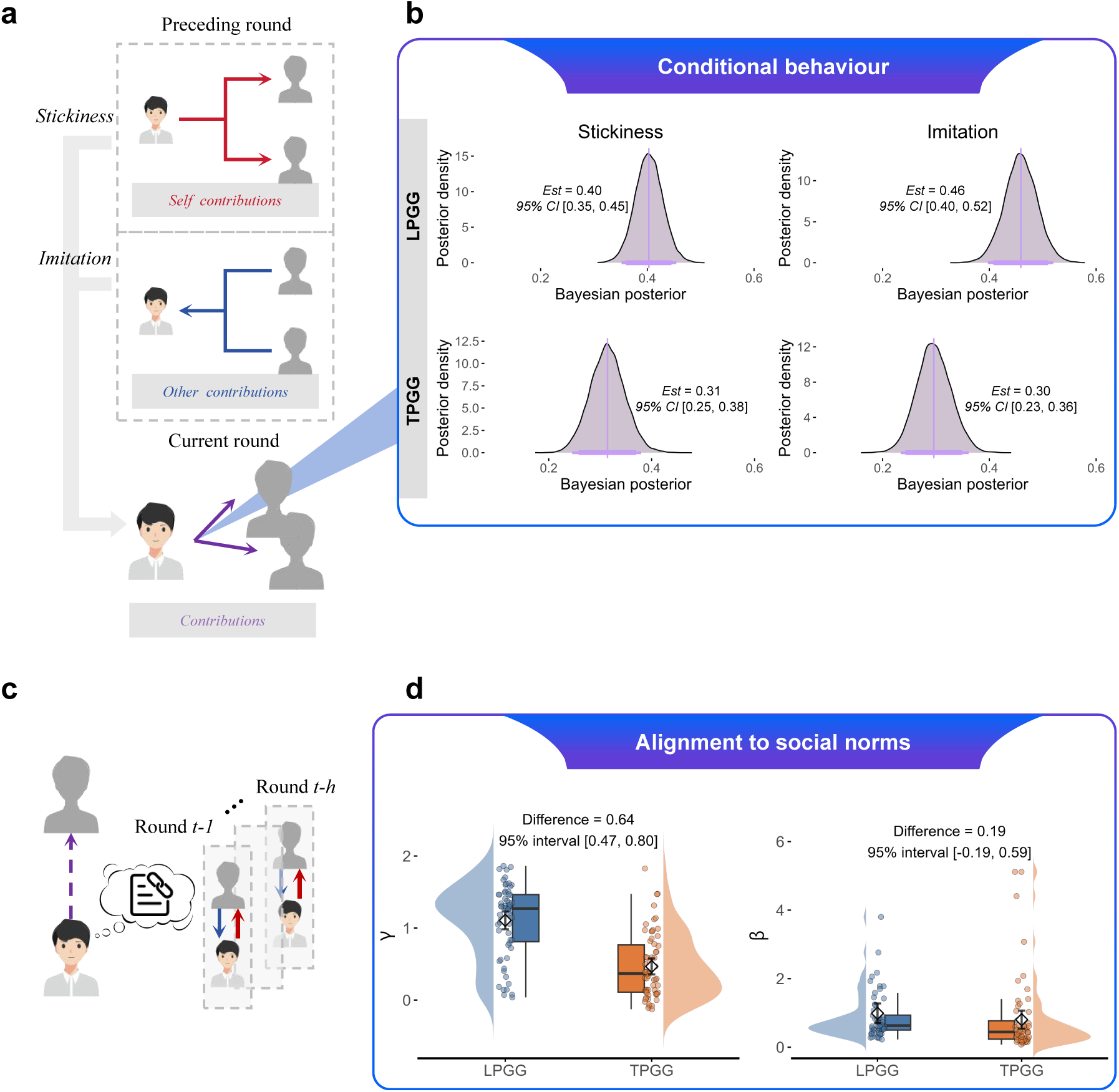
Conditional behavior and norm–utility modelling. **a,** Schematic of the conditional-contribution model. Contributions to a given dyadic public pool in the current round were modelled as a function of the focal individual’s and their partner’s contributions to the same pool in the preceding round, corresponding to stickiness and imitation, respectively [43–45]. **b,** Posterior distributions of the Bayesian regression coefficients, with 90% and 95% highest-density intervals. Both stickiness and imitation were stronger in the LPGG than in the TPGG. **c,** Schematic of the norm–utility model [39, 40]. For each decision, the model evaluated all feasible allocations between the two partners and assigned each option a subjective utility combining its material payoff with the cost of deviating from partner-specific, history-dependent contribution standards. **d,** Individual-level posterior estimates of the normative preference weight, *γ*, and the inverse-temperature parameter, *β*. Each point represents one participant. The weight *γ* quantifies the extent to which contributions align with dyadic historical standards, whereas *β* captures the sensitivity of decisions to differences in utility.

**Fig. 3:**
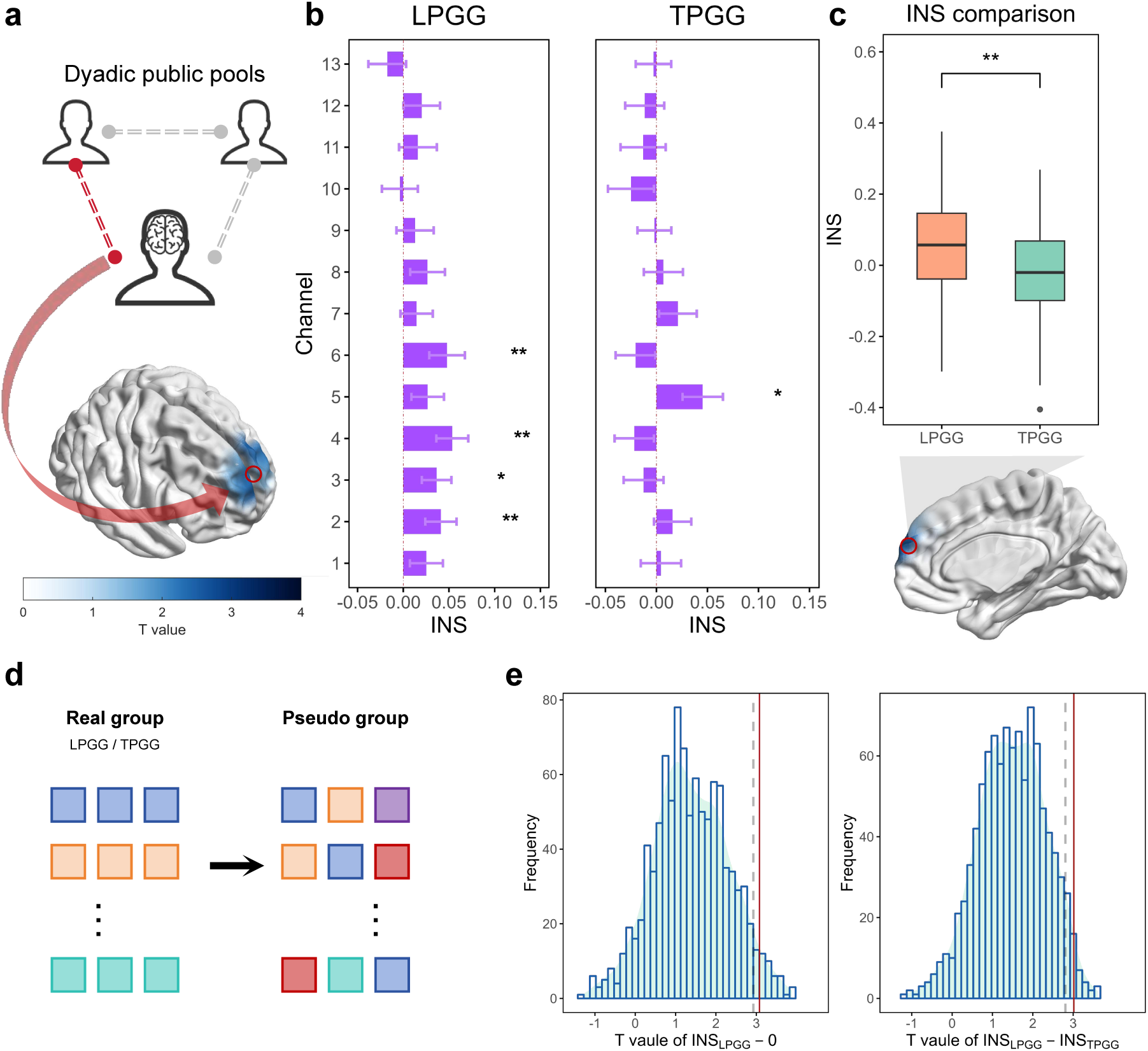
Dyadic inter-brain neural synchrony. **a,** Quantification of dyadic inter-brain neural synchrony (INS) and its spatial distribution over the anterior prefrontal cortex in the LPGG. Wavelet transform coherence (WTC) was calculated for each pair of group members, corrected for resting-state coherence and averaged across rounds to obtain dyadic INS. **b,** Channel-wise task-related INS in the LPGG and TPGG. Channels 2, 3, 4 and 6 showed significant increases in the LPGG after FDR. **c,** Dyadic INS in Channel 4 across game structures. Synchrony was significantly stronger in the LPGG than in the TPGG. **d,** Construction of pseudo-groups. Participants were reassigned within each game structure under the constraint that no pseudo-group contained more than one member of the same real interacting group. **e,** Pseudo-group permutation results for Channel 4 based on 1,000 permutations. Channel-4 INS in real LPGG groups exceeded the corresponding pseudo-group distribution and the observed LPGG–TPGG difference was greater than expected under pseudo-group reassignment.

Bayesian analyses yielded higher estimates of imitation in the LPGG than in the TPGG (Fig. 2b, right panel; LPGG: *β*_imitation_ = 0.46, SD = 0.03, 95% credible interval [0.40, 0.52], *R̂* = 1.00; TPGG: *β*_imitation_ = 0.30, SD = 0.03, 95% credible interval [0.23, 0.36], *R̂* = 1.00). Estimates of stickiness were also higher in the LPGG (Fig. 2b, left panel; LPGG: *β*_stickiness_ = 0.40, SD = 0.03, 95% credible interval [0.35, 0.45], *R̂* = 1.00; TPGG: *β*_stickiness_ = 0.31, SD = 0.03, 95% credible interval [0.25, 0.38], *R̂* = 1.00). Additional analyses supported the robustness of this pattern (Supplementary Fig. 5). Together, these estimates suggest that individuals in the LPGG showed greater stickiness to their own contributions alongside greater responsiveness to their partners’ previous contributions. Under free allocation, this combination is consistent with selective reciprocity, whereby contributions to a given pool are adjusted in response to contributions previously received from the same partner.

The preceding analyses characterized contribution dynamics and conditional behavior, but did not directly quantify the role of social norms under free allocation. We therefore fitted a Bayesian softmax choice model to estimate a normative preference weight, *γ* [39, 40, 46]. For each dyadic pool, we defined a history-dependent contribution standard, *c*^∗^, as the mean of the two members’ previous contributions to that pool. The model distinguished sensitivity to material payoffs from sensitivity to deviations from this shared contribution standard. Higher values of *γ* indicate stronger alignment to history-dependent contribution standards, whereas lower values indicate weaker alignment. We compared preference weight (*γ*) and inverse temperature (*β*) between game structures by independently resampling individual-level posterior means. *γ* was higher in the LPGG than in the TPGG (Fig. 2d; LPGG: *γ* = 1.107; TPGG: *γ* = 0.467; 95% bootstrap interval for the LPGG–TPGG difference [0.471, 0.803]), indicating stronger alignment to history-dependent contribution standards. By contrast, the corresponding interval for inverse temperature included zero (LPGG: *β* = 0.989; TPGG: *β* = 0.803; 95% bootstrap interval [−0.188, 0.586]), providing no clear evidence of a difference in how strongly choices favored higher-utility options.

### The LPGG with free allocation elicited stronger inter-brain neural synchrony

Having characterized distinct behavioral dynamics and norms under free allocation in the two game structures, we next asked whether these differences were accompanied by distinct neural responses during interaction. We analyzed functional near-infrared spectroscopy (fNIRS) time series of oxygenated haemoglobin concentration to characterize both dyadic inter-brain synchrony and individual neural activity. We first examined inter-brain synchrony as a potential neural correlate of interpersonal coordination and alignment with shared contribution standards [27, 47]. As free allocation allowed individuals to adjust their contributions to specific partners within each public pool, we quantified pairwise neural coupling using wavelet transform coherence between partners’ fNIRS signals. Inter-brain neural synchrony (INS) was defined as coupling during the operation phase, corrected for resting-state coupling between the same two individuals.

We first assessed task-related INS within each game structure. In the LPGG, INS increased significantly relative to rest in anterior prefrontal channels 2, 3, 4 and 6 (Fig. 3a, b; Channel 2: *t*(59) = 2.41, *P* = 0.0191, *P*_FDR_ = 0.0446; channel 3: *t*(59) = 2.26, *P* = 0.0278, *P*_FDR_ = 0.0486; channel 4: *t*(59) = 3.08, *P* = 0.00315, *P*_FDR_ = 0.0220; channel 6: *t*(59) = 2.46, *P* = 0.0168, *P*_FDR_ = 0.0446). In the TPGG, an increase was only observed in dmPFC channel 5, but did not survive correction for multiple comparisons (Fig. 3b; Channel 5: *t*(59) = 2.30, *P* = 0.0253, *P*_FDR_ = 0.177). Between-structure comparisons revealed higher mean INS in the anterior dmPFC in the LPGG than in the TPGG, with a significant difference at channel 4 (*β* = 0.075, SE = 0.026, *t*(118) = 2.90, *P* = 0.00442). Analyses using 1,000 pseudo-group permutations further supported this finding: channel-4 INS in the LPGG exceeded that expected from pseudo-groups (*P* = 0.038), and the LPGG–TPGG difference was greater for real interacting groups than for pseudo-groups (*P* = 0.020). Having identified stronger task-related INS in the LPGG, we next examined its relationship with dyadic behavior. In the LPGG, INS at dmPFC channel 3 was negatively associated with norms, indexed by the mean *γ* of the two partners (Fig. 4a; *β* = −0.103, SE = 0.036, *t*(58) = −2.88, *P* = 0.00550). Thus, stronger neural coupling was associated with weaker adherence to established dyadic contribution standards. We then examined whether the same INS measure was associated with contribution differences between partners. Dyads with larger contribution differences averaged across rounds showed stronger channel-3 INS (Fig. 4b; *β* = 0.016, SE = 0.008, *t*(58) = 2.10, *P* = 0.0404). Sliding-window analyses spanning two to six rounds revealed no consistent phase-specific association after FDR correction. Associations were instead observed in cumulative windows, suggesting that the relationship was more apparent when contribution differences were aggregated over longer interaction histories [48, 49] (Supplementary Fig. 8). Together, these findings are consistent with dmPFC synchrony reflecting ongoing interpersonal coordination in the free-allocation LPGG, potentially involving the monitoring and adjustment of partner-specific reciprocal strategies, rather than stronger adherence to established contribution standards.

**Fig. 4:**
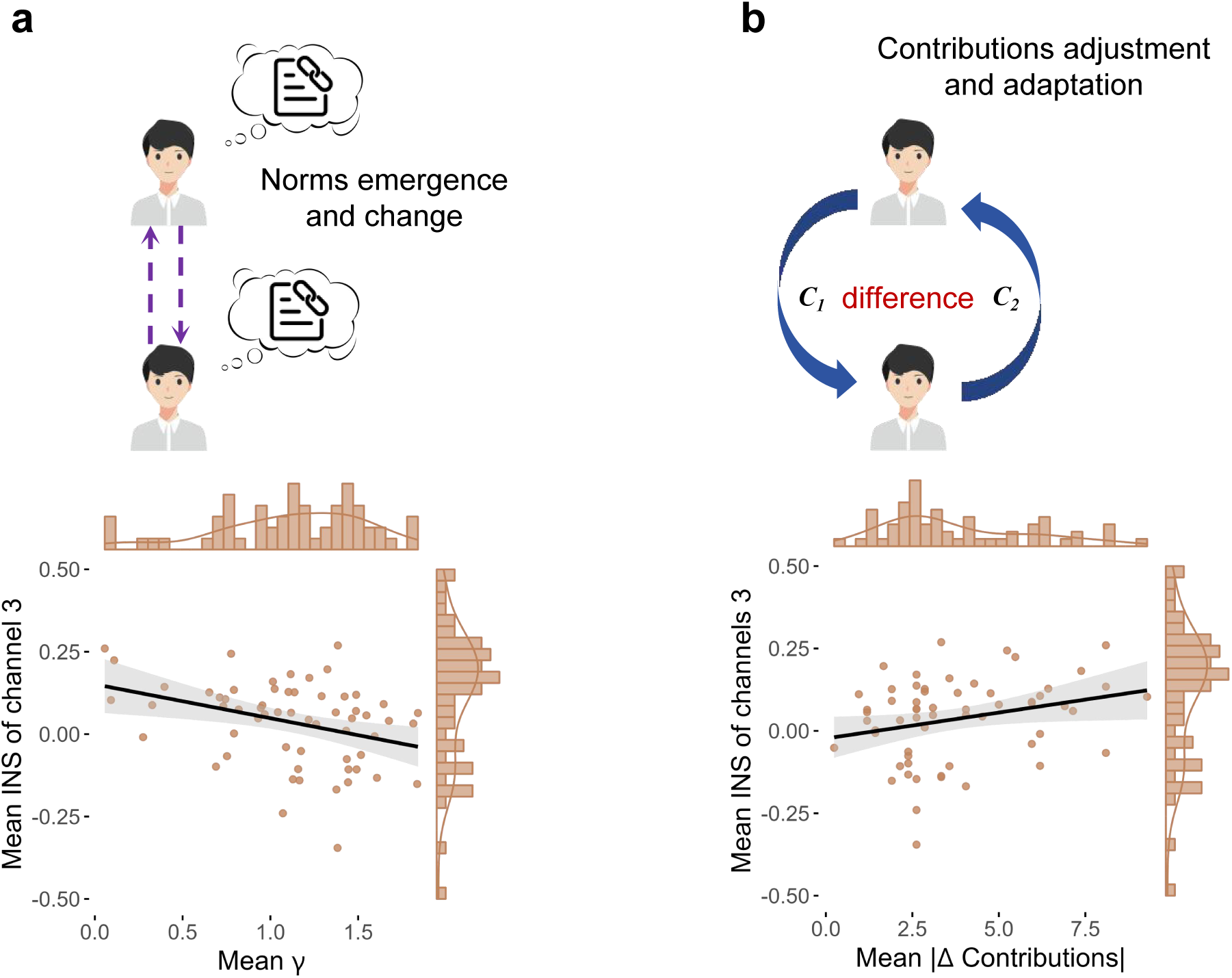
Associations between inter-brain neural synchrony and behavioral dynamics in the LPGG. **a,** Dyadic mean normative preference weight was negatively associated with interbrain neural synchrony (INS) in Channel 3. **b,** Mean partner-specific contribution differences were positively associated with Channel-3 INS.

**Fig. 5:**
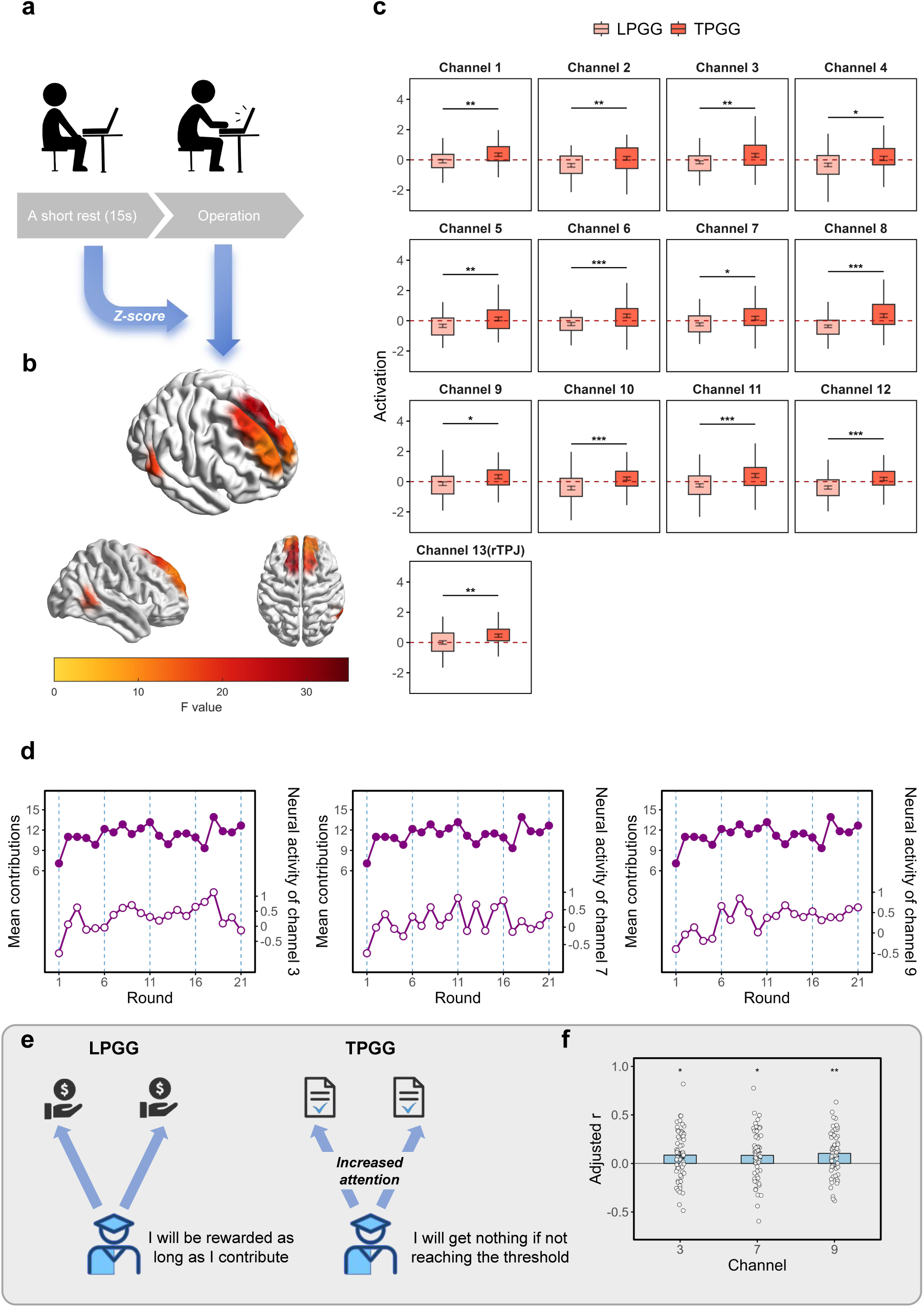
Individual neural activity and its association with contribution behavior. **a,** Quantification of task-related neural activity. The 15-s period preceding the operation phase served as the baseline, and oxygenated haemoglobin signals during the operation phase were standardized using the corresponding baseline mean and standard deviation. **b,c,** Differences in participant-level neural activity between the LPGG and TPGG. Activity was generally stronger in the TPGG than in the LPGG. **d,** Round-level associations between mean total contributions and neural activity during the operation phase in the TPGG. Activity in Channels 3, 7 and 9 was positively associated with mean total contributions. **e,** Schematic illustration of the distinct decisions and threshold-related risks in the LPGG and TPGG. **f,** Within-participant associations between round-by-round neural activity and total contributions. In the TPGG, greater activity in Channels 3, 7 and 9 was associated with higher individual contributions in the corresponding round.

### Neural signatures reflect individual risk evaluation and calculated greed

We next examined whether individual neural responses differed between game structures under free allocation. We first compared oxygenated haemoglobin (oxy-Hb) concentrations during the operation phase (Fig. 5a). Responses were generally greater in the TPGG than in the LPGG, including across several channels over the dmPFC and rTPJ (Fig. 5b, c).

We then examined the relationship between neural activity and contributions at both the round and individual levels. To characterize behavioral dynamics, we first averaged individuals’ total contributions within each round and related these values to the corresponding mean neural activity. In the TPGG, activity in dmPFC channels 3, 7 and 9 was positively associated with mean round-level contributions (Fig. 5d; Channel 3: *β* = 0.150, SE = 0.057, *t*(19) = 2.61, *P* = 0.0172; Channel 7: *β* = 0.151, SE = 0.048, *t*(19) = 3.12, *P* = 0.00571; Channel 9: *β* = 0.121, SE = 0.042, *t*(19) = 2.90, *P* = 0.00924). We next examined whether these associations were also evident at the individual level. Greater activity in each of the three channels was associated with higher individual contributions (Fig. 5e, f; Channel 3: *t*(56) = 2.48, *P* = 0.0160, 95% CI [0.017, 0.155]; Channel 7: *t*(56) = 2.44, *P* = 0.0177, 95% CI [0.015, 0.153]; Channel 9: *t*(56) = 3.43, *P* = 0.00115, 95% CI [0.043, 0.165]). In the LPGG, although activity in several channels was associated with mean round-level contributions, none showed a statistically significant association with individual contributions.

One possible interpretation is that these neural responses reflect individual evaluations of risk and uncertainty surrounding threshold attainment, rather than alignment to group-level social norms. In the TPGG, the expected return per unit contributed was lower and depended strongly on the part-ner’s contribution. Under free allocation, individuals also had to assess which public pool was more likely to reach the threshold. Larger contributions thus committed more resources to an uncertain outcome, potentially increasing pressure and cognitive demands [50]. In the LPGG, by contrast, contributions generated returns regardless of threshold attainment, reducing this source of uncertainty (Fig. 5e).

We also quantified within-brain functional connectivity using coherence across 12 channel pairs linking the dmPFC and rTPJ. Following the neural activity analyses, we examined associations between connectivity and individual contributions across rounds, testing Fisher-*z*-transformed within-individual correlations against zero. In the LPGG, stronger coherence between dmPFC channels 1, 3, 5, 7 and 12 and rTPJ channel 13 was associated with lower contributions (Fig. 6; Channel 1–13: *t*(57) = −2.03, *P* = 0.0469, 95% CI [−0.160, −0.001]; Channel 3–13: *t*(58) = −2.11, *P* = 0.0394, 95% CI [−0.160, −0.004]; Channel 5–13: *t*(58) = −2.29, *P* = 0.0256, 95% CI [−0.146, −0.010]; Channel 7–13: *t*(58) = −2.18, *P* = 0.0331, 95% CI [−0.134, −0.006]; Channel 12–13: *t*(58) = −2.09, *P* = 0.0412, 95% CI [−0.144, −0.003]). In the TPGG, no channel pair showed a statistically significant association with individual contributions across rounds. The pattern suggests that dmPFC–rTPJ connectivity in the LPGG may reflect processes involved in evaluating personal payoffs relative to the benefits of contribution. In LPGG, where individual payoff maximization favors lower contribution, such deliberation may shift behavior toward calculated self-interest rather than intuitive contribution, consistent with the “spontaneous giving and calculated greed” account [51].

**Fig. 6:**
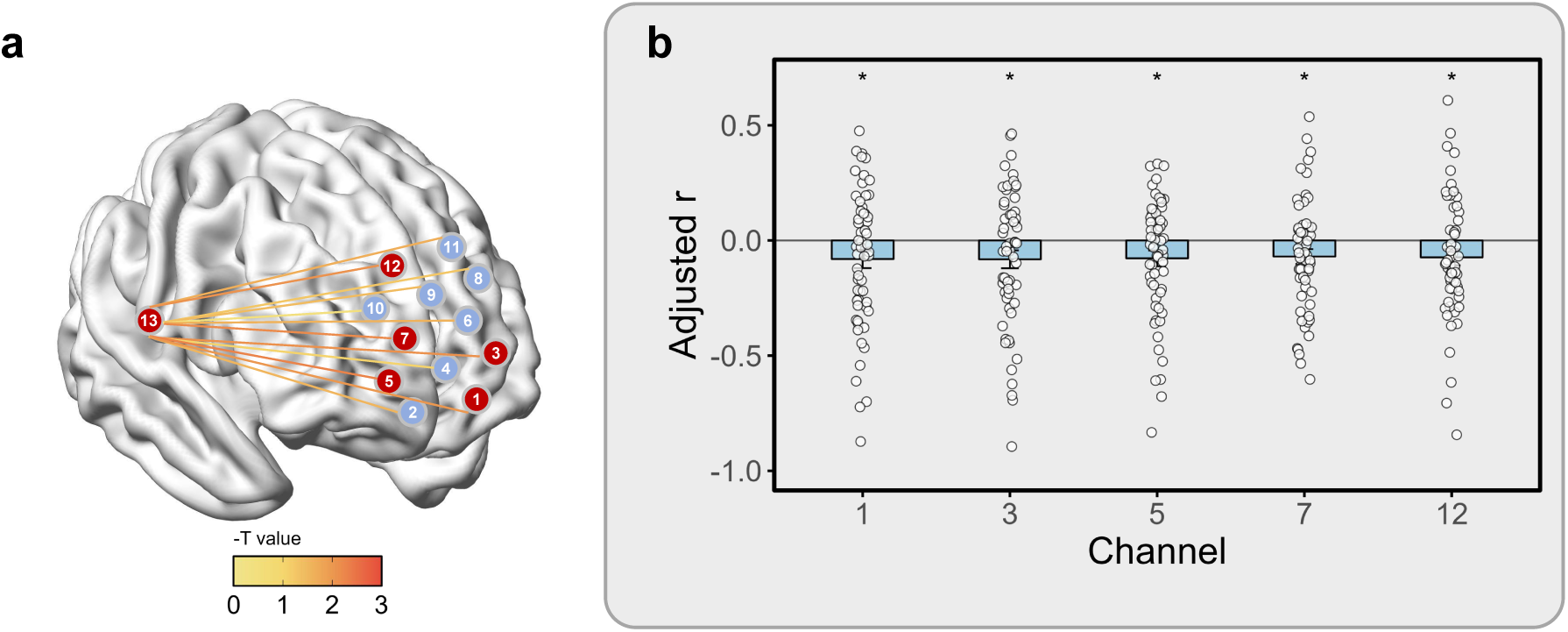
Associations between intra-brain functional connectivity and individual contributions. **a,b,** Within-participant associations between round-by-round dmPFC–rTPJ functional connectivity and total contributions. In the LPGG, stronger connectivity between rTPJ Channel 13 and dmPFC Channels 1, 3, 5, 7 and 12 was associated with lower individual contributions in the corresponding round.

Together, these findings characterize individual-level neural responses associated with contribution decisions under free allocation in the two game structures. In the TPGG, greater dmPFC activity was associated with higher contributions, consistent with a possible role for evaluating uncertain outcomes surrounding threshold attainment. In the LPGG, stronger dmPFC–rTPJ connectivity was associated with lower contributions, suggesting a potential role for interactions between these regions in contribution decisions.

## Discussion

Our study examines how game structure relates to the behavioral and neural signatures of cooperation in a free-allocation setting by integrating analyses of behavioral dynamics with fNIRS hyperscanning. Contrary to the theoretical expectation that threshold incentives facilitate norm-based coordination, we did not observe stronger alignment to history-dependent contribution standards in the TPGG. Instead, the LPGG was characterized by greater alignment to these standards, stronger partner-specific reciprocal adjustment, and enhanced dyadic inter-brain synchrony in the dmPFC. The results suggest that, in real-world social interactions, the motives to cooperate depend not only on the game structures, but also on whether individuals can freely allocate effort across multiple social partners. When cooperation is embedded in networked relationships [52, 53], the individuals may track not simply whether contributions meet the expectations, but how reciprocal relationships are identified, compared and updated across different partners [16, 37, 54, 55].

Existing theory identifies the introduction of a threshold as a qualitative change in the incentive structure of a linear social dilemma. Under standard provision-point designs, TPGGs can admit positive-contribution, Pareto-efficient Nash equilibria, turning the cooperation problem into one of coordination around a salient collective target [2, 56]. This theoretical perspective suggests that, relative to the LPGG, the TPGG may promote convergence on shared contribution standards and thereby strengthen norm alignment [2, 9, 57]. Our findings suggest, however, that this expectation need not hold under free allocation. In the LPGG, individuals increasingly differentiated their contributions between partners over successive rounds, consistent with more selective allocation within dyadic relationships. No comparable temporal trend was detected in the TPGG. Moreover, the norm-utility model yielded substantially higher normative preference weight, *γ*, in the LPGG, indicating stronger sensitivity to history-dependent contribution standards within each dyadic pool [40, 46]. These findings suggest that cooperation under free allocation is characterized not only by how much individuals contribute, but also by how selectively they allocate contributions and how closely their decisions align with the standards established within particular relationships.

The hyperscanning findings provided a neural perspective on these behavioral patterns. In the LPGG, task-related dyadic inter-brain synchrony increased significantly in anterior dmPFC Channels 2, 3, 4 and 6, with Channel 4 showing stronger synchrony than in the TPGG. Pseudo-group permutation tests further supported the specificity of this effect to real interacting partners [58]. Given the involvement of medial prefrontal systems in social inference, monitoring others’ intentions and processing socially relevant information [30, 31, 33, 34], enhanced synchrony may reflect the interpersonal demands of maintaining and adjusting partner-specific contribution strategies. This interpretation is consistent with accounts linking inter-brain synchrony to coordinated social interaction [23, 24]. Importantly, within the LPGG, stronger INS at dmPFC Channel 3 was associated with lower dyadic mean preference weight, *γ*, and larger contribution differences between partners. These associations argue against a simple interpretation of synchrony as stronger alignment to social norms or greater behavioral convergence. Instead, they are consistent with the possibility that dmPFC synchrony reflects the coordination demands arising when partners differ in their contributions and adjust their reciprocal strategies over repeated interaction.

Although the TPGG did not exhibit stronger dyadic INS, it elicited greater individual neural activity, with dmPFC Channels 3, 7 and 9 showing positive associations with contributions at both the round and individual levels. This pattern suggests that weaker inter-brain synchrony need not imply weaker individual neural engagement. Because returns in the TPGG depended on joint contributions reaching a threshold, contribution decisions involved uncertainty about partners’ behavior and the likelihood of reaching the threshold [2]. The positive activity–contribution associations may therefore reflect processes involved in predicting partners’ contributions and evaluating uncertain outcomes [59–62]. In the LPGG, stronger dmPFC–rTPJ connectivity was instead associated with lower individual contributions. Given the private incentive to withhold contributions when the marginal per-capita return is below one, this pattern is compatible with greater consideration of personal costs and payoffs and therefore interpreted as calculated greed [51].

Taken together, these neural findings suggest that free allocation may place different demands on interpersonal coordination and individual decision across game structures. In the LPGG, enhanced dyadic INS was consistent with greater demands for partner-specific monitoring and reciprocal adjustment, rather than stronger alignment to established social norms. Also, exploratory associations between dmPFC–rTPJ connectivity and lower contributions may reflect consideration of private costs and payoffs. In the TPGG, threshold-dependent payoffs may increase uncertainty about each partner’s contribution and the likelihood of successful provision, potentially weakening stickiness to history-dependent contribution standards. Greater individual dmPFC activity in this condition is compatible with these predictive and evaluative demands. This account suggests a potential boundary condition for the coordination advantage of threshold incentives: under free allocation, an explicit collective target need not produce stronger norm alignment and may instead place greater demands on evaluating uncertain outcomes [12].

Real-world cooperation often requires individuals to allocate limited time, attention and resources across multiple tasks, projects and partners [63, 64]. Employees balance competing responsibilities, researchers divide effort among collaborations. In these settings, introducing a threshold or performance target may not be sufficient to strengthen alignment to social norms. Under free allocation, thresholds may shift attention towards the likelihood of successful provision and the risk of unre-warded contributions, potentially reducing reliance on partner-specific interaction histories [65]. By contrast, continuous returns and feedback that can be attributed to specific partners may help individuals interpret reciprocal behavior and develop contribution standards [54]. Our findings therefore suggest that cooperative institutions should consider how performance targets interact with allocation freedom [53].

Together, our findings suggest that the relationship between game structure and cooperation depends on how individuals allocate resources across partners. Under free allocation, the LPGG was associated with greater partner-specific differentiation, stronger sensitivity to history-dependent contribution standards and enhanced dyadic INS, whereas the TPGG elicited greater individual neural activity, potentially reflecting demands associated with uncertain threshold attainment. These patterns suggest that an explicit collective target need not translate into stronger norm alignment when contributions are distributed across multiple pools. They also refine the interpretation of INS: stronger coupling may reflect ongoing reciprocal adjustment rather than an established consensus. By linking contribution dynamics, computational estimates of norms and neural responses, this study highlights the importance of partner-specific interactions in understanding how cooperative standards emerge and guide behavior [66].

## Supporting information

Supplementary Methods and Results

## Data availability

Behavioral data and processed multidimensional functional near-infrared spectroscopy (fNIRS) data from the four-person groups are publicly available on the Open Science Framework (OSF) at (https://osf.io/gm54w/).

## Code availability

The code used to reproduce the statistical analyses is publicly available on the Open Science Framework (OSF) at (https://osf.io/gm54w/).

## Acknowledgements

This work is supported by National Cyber Security-National Science and Technology Major Project (2025ZD1503700), Program of National Natural Science Foundation of China (12425114, 12671643, 12501702, 12526528, 62441617), Taishan Scholars Program of Shandong Province (tstp20230635), Key R&D Program of Shandong Province (2026CXPT283), the Fundamental Research Funds for the Central Universities, the Opening Project of the State Key Laboratory of General Artificial Intelligence(Project NSKLAGI2025OP16), the special Funding for the ‘Case-by-Case-Introduction of Top Talent (Teams)’, and Beijing Advanced Innovation Center for Future Blockchain and Privacy Computing.

## Author contributions

Y.L. and W.X. conceived the project and designed the experiments. Y.L., W.X., Y.W. and W.D. organized the experiments and collected the data. Y.L., X.W., W.D. and L.L. analyzed the data and prepared the figures. Y.L. developed the computational model and interpreted the results. Y.L., W.X., Y.W., X.W., X.C., L.L. and S.T. wrote the original and final version of the manuscript. H.Z. and S.T. funded and supervised the project.

## Competing interests

The authors declare no competing interests.

**Correspondence and requests for materials** should be addressed to Shaoting Tang.

## Notes

### Competing Interest Statement

The authors have declared no competing interest.

## References

[1] Jin, S., Spadaro, G. & Balliet, D. Institutions and cooperation: A meta-analysis of structural features in social dilemmas. Journal of personality and social psychology 129, 286 (2025).

[2] Abele, S., Stasser, G. & Chartier, C. Conflict and coordination in the provision of public goods: A conceptual analysis of continuous and step-level games. Personality and Social Psychology Review 14, 385–401 (2010).

[3] Buchholz, W. & Sandler, T. Global public goods: a survey. Journal of Economic Literature 59, 488–545 (2021).

[4] Balliet, D. & Lindström, B. Inferences about interdependence shape cooperation. Trends in Cognitive Sciences 27, 583–595 (2023).

[5] Fehr, E. & Fischbacher, U. Social norms and human cooperation. Trends in cognitive sciences 8, 185–190 (2004).

[6] Tu, C., D’Odorico, P., Li, Z. & Suweis, S. The emergence of cooperation from shared goals in the governance of common-pool resources. Nature Sustainability 6, 139–147 (2023).

[7] Horne, C. & Mollborn, S. Norms: An integrated framework. Annual Review of Sociology 46, 467–487 (2020).

[8] Ostrom, E. Collective action and the evolution of social norms. Journal of economic perspectives 14, 137–158 (2000).

[9] Szekely, A. et al. Evidence from a long-term experiment that collective risks change social norms and promote cooperation. Nature communications 12, 5452 (2021).

[10] Wang, J., Fu, F., Wu, T. & Wang, L. Emergence of social cooperation in threshold public goods games with collective risk. *Physical Review E—Statistical*, Nonlinear, and Soft Matter Physics 80, 016101 (2009).

[11] Columbus, S. & Molho, C. Subjective interdependence and prosocial behaviour. Current Opinion in Psychology 43, 226–231 (2022).

[12] Deutchman, P., Amir, D., Jordan, M. R. & McAuliffe, K. Common knowledge promotes cooperation in the threshold public goods game by reducing uncertainty. Evolution and Human Behavior 43, 155–167 (2022).

[13] Santos, F. C., Santos, M. D. & Pacheco, J. M. Social diversity promotes the emergence of cooperation in public goods games. Nature 454, 213–216 (2008).

[14] Bikard, M., Murray, F. & Gans, J. S. Exploring trade-offs in the organization of scientific work: Collaboration and scientific reward. Management science 61, 1473–1495 (2015).

[15] Bak-Coleman, J. B. et al. Stewardship of global collective behavior. Proceedings of the National Academy of Sciences 118, e2025764118 (2021).

[16] Melis, A. P. & Raihani, N. J. The cognitive challenges of cooperation in human and nonhuman animals. Nature Reviews Psychology 2, 523–536 (2023).

[17] Kingsbury, L. & Hong, W. A multi-brain framework for social interaction. Trends in neurosciences 43, 651–666 (2020).

[18] Park, S. A., Sestito, M., Boorman, E. D. & Dreher, J.-C. Neural computations underlying strategic social decision-making in groups. Nature communications 10, 5287 (2019).

[19] Yoshida, W., Seymour, B., Friston, K. J. & Dolan, R. J. Neural mechanisms of belief inference during cooperative games. Journal of Neuroscience 30, 10744–10751 (2010).

[20] Fehr, E. & Schurtenberger, I. Normative foundations of human cooperation. Nature human behaviour 2, 458–468 (2018).

[21] Réveillé, C., Vergotte, G., Perrey, S. & Bosselut, G. Using interbrain synchrony to study teamwork: a systematic review and meta-analysis. Neuroscience & Biobehavioral Reviews 159, 105593 (2024).

[22] Xie, H. et al. Finding the neural correlates of collaboration using a three-person fmri hyperscanning paradigm. Proceedings of the National Academy of Sciences 117, 23066–23072 (2020).

[23] Gordon, I. & Bartsch, R. P. Correlates of interpersonal physiological synchrony and sources of empirical heterogeneity. Nature Reviews Psychology 1–15 (2026).

[24] Chidichimo, E. et al. Towards an informational account of interpersonal coordination. Nature Reviews Neuroscience 27, 121–137 (2026).

[25] Cui, X., Bryant, D. M. & Reiss, A. L. Nirs-based hyperscanning reveals increased interpersonal coherence in superior frontal cortex during cooperation. Neuroimage 59, 2430–2437 (2012).

[26] Hasson, U. & Frith, C. D. Mirroring and beyond: coupled dynamics as a generalized framework for modelling social interactions. Philosophical Transactions of the Royal Society B: Biological Sciences 371, 20150366 (2016).

[27] Schilbach, L. & Redcay, E. Synchrony across brains. Annual review of psychology 76, 883–911 (2025).

[28] Hakim, U. et al. Quantification of inter-brain coupling: A review of current methods used in haemodynamic and electrophysiological hyperscanning studies. NeuroImage 280, 120354 (2023).

[29] Zoh, Y., Chang, S. W. & Crockett, M. J. The prefrontal cortex and (uniquely) human cooperation: a comparative perspective. Neuropsychopharmacology 47, 119–133 (2022).

[30] Buckholtz, J. W. & Marois, R. The roots of modern justice: cognitive and neural foundations of social norms and their enforcement. Nature neuroscience 15, 655–661 (2012).

[31] Olsson, A., Knapska, E. & Lindström, B. The neural and computational systems of social learning. Nature Reviews Neuroscience 21, 197–212 (2020).

[32] Ruff, C. C. & Fehr, E. The neurobiology of rewards and values in social decision making. Nature Reviews Neuroscience 15, 549–562 (2014).

[33] Ruff, C. C., Ugazio, G. & Fehr, E. Changing social norm compliance with noninvasive brain stimulation. Science 342, 482–484 (2013).

[34] Suzuki, S., Adachi, R., Dunne, S., Bossaerts, P. & O’Doherty, J. P. Neural mechanisms underlying human consensus decision-making. Neuron 86, 591–602 (2015).

[35] Wolff, A. & Dumas, G. Coupled neural timescales in social interaction. Trends in Cognitive Sciences (2026).

[36] Jia, D. et al. Social networking agency and prosociality are inextricably linked in economic games. Nature Human Behaviour 9, 2620–2631 (2025).

[37] Rossetti, C. S., Hauser, O. P. & Hilbe, C. Dynamics of cooperation in concurrent games. Nature Communications 16, 1524 (2025).

[38] Babitz, D. & Eldar, E. How social norms emerge: The interindividual actor–critic. Psychological review (2025).

[39] Brock, W. A. & Durlauf, S. N. Discrete choice with social interactions. The Review of Economic Studies 68, 235–260 (2001).

[40] Gavrilets, S., Tverskoi, D. & Sánchez, A. Modelling social norms: an integration of the norm-utility approach with beliefs dynamics. Philosophical Transactions of the Royal Society B 379, 20230027 (2024).

[41] Barrett, S. & Dannenberg, A. Sensitivity of collective action to uncertainty about climate tipping points. Nature Climate Change 4, 36–39 (2014).

[42] Gelfand, M. J., Gavrilets, S. & Nunn, N. Norm dynamics: Interdisciplinary perspectives on social norm emergence, persistence, and change. Annual Review of Psychology 75, 341–378 (2024).

[43] Dong, Y., Zhang, B. & Tao, Y. The dynamics of human behavior in the public goods game with institutional incentives. Scientific Reports 6, 28809 (2016).

[44] Press, W. H. & Dyson, F. J. Iterated prisoner’s dilemma contains strategies that dominate any evolutionary opponent. Proceedings of the National Academy of Sciences 109, 10409–10413 (2012).

[45] Pisauro, M. A., Fouragnan, E., Arabadzhiyska, D., Apps, M. A. & Philiastides, M. G. Neural implementation of computational mechanisms underlying the continuous trade-off between cooperation and competition. Nature Communications 13, 6873 (2022).

[46] Alger, I., Gavrilets, S. & Durkee, P. Proximate and ultimate drivers of norms and norm change. Current Opinion in Psychology 60, 101916 (2024).

[47] Shamay-Tsoory, S. G., Saporta, N., Marton-Alper, I. Z. & Gvirts, H. Z. Herding brains: a core neural mechanism for social alignment. Trends in cognitive sciences 23, 174–186 (2019).

[48] Gherman, S. et al. Intracranial electroencephalography reveals effector-independent evidence accumulation dynamics in multiple human brain regions. Nature Human Behaviour 8, 758–770 (2024).

[49] Pesaran, B. et al. Investigating large-scale brain dynamics using field potential recordings: analysis and interpretation. Nature neuroscience 21, 903–919 (2018).

[50] Cocchi, L., Zalesky, A., Fornito, A. & Mattingley, J. B. Dynamic cooperation and competition between brain systems during cognitive control. Trends in cognitive sciences 17, 493–501 (2013).

[51] Rand, D. G., Greene, J. D. & Nowak, M. A. Spontaneous giving and calculated greed. Nature 489, 427–430 (2012).

[52] Melamed, D., Harrell, A. & Simpson, B. Cooperation, clustering, and assortative mixing in dynamic networks. Proceedings of the National Academy of Sciences 115, 951–956 (2018).

[53] Rand, D. G., Arbesman, S. & Christakis, N. A. Dynamic social networks promote cooperation in experiments with humans. Proceedings of the National Academy of Sciences 108, 19193–19198 (2011).

[54] Van Lange, P. A. & Bergmann, P. K. Human cooperation is strong among individuals but fragile between groups. Nature Reviews Psychology 1–14 (2026).

[55] Hilbe, C., Chatterjee, K. & Nowak, M. A. Partners and rivals in direct reciprocity. Nature human behaviour 2, 469–477 (2018).

[56] Bagnoli, M. & Lipman, B. L. Provision of public goods: Fully implementing the core through private contributions. The Review of Economic Studies 56, 583–601 (1989).

[57] Tavoni, A., Dannenberg, A., Kallis, G. & Löschel, A. Inequality, communication, and the avoidance of disastrous climate change in a public goods game. Proceedings of the National Academy of Sciences 108, 11825–11829 (2011).

[58] Koul, A., Ahmar, D., Iannetti, G. D. & Novembre, G. Spontaneous dyadic behavior predicts the emergence of interpersonal neural synchrony. NeuroImage 277, 120233 (2023).

[59] Konovalov, A., Hill, C., Daunizeau, J. & Ruff, C. C. Dissecting functional contributions of the social brain to strategic behavior. Neuron 109, 3323–3337 (2021).

[60] Hill, C. A. et al. A causal account of the brain network computations underlying strategic social behavior. Nature neuroscience 20, 1142–1149 (2017).

[61] Redcay, E. & Schilbach, L. Using second-person neuroscience to elucidate the mechanisms of social interaction. Nature Reviews. Neuroscience 20, 495 (2019).

[62] Bach, D. R. & Dolan, R. J. Knowing how much you don’t know: a neural organization of uncertainty estimates. Nature reviews neuroscience 13, 572–586 (2012).

[63] Battiston, F. et al. Higher-order interactions shape collective human behaviour. Nature Human Behaviour 1–17 (2025).

[64] Allen, B. Understanding large-scale cooperation in a nonlinear, interconnected world. Nature Computational Science 6, 112–114 (2026).

[65] Cárdenas, J.-C. et al. Fragility of the provision of local public goods to private and collective risks. Proceedings of the National Academy of Sciences 114, 921–925 (2017).

[66] Gvirts, H. Z. & Perlmutter, R. What guides us to neurally and behaviorally align with anyone specific? a neurobiological model based on fnirs hyperscanning studies. The Neuroscientist 26, 108–116 (2020).

[67] Falk, A., Becker, A., Dohmen, T., Huffman, D. & Sunde, U. The preference survey module: A validated instrument for measuring risk, time, and social preferences. Management Science 69, 1935–1950 (2023).

[68] Murphy, R. O., Ackermann, K. A. & Handgraaf, M. J. Measuring social value orientation. Judgment and Decision making 6, 771–781 (2011).

[69] Wang, G., Li, J., Wang, W., Niu, X. & Wang, Y. Confusion cannot explain cooperative behavior in public goods games. Proceedings of the National Academy of Sciences 121, e2310109121 (2024).

[70] Zimeo Morais, G. A., Balardin, J. B. & Sato, J. R. fnirs optodes’ location decider (fold): a toolbox for probe arrangement guided by brain regions-of-interest. Scientific reports 8, 3341 (2018).

[71] Ye, J. C., Tak, S., Jang, K. E., Jung, J. & Jang, J. Nirs-spm: statistical parametric mapping for near-infrared spectroscopy. Neuroimage 44, 428–447 (2009).

[72] Bürkner, P.-C. brms: An r package for bayesian multilevel models using stan. Journal of statistical software 80, 1–28 (2017).

