## Supplementary Methods and Results for "Game structure shapes normative preferences and inter-brain neural synchrony in free-allocation public goods games"

**5 Pre-experiment questionnaire** **18**

**6 Post-experiment measure** **20**

**7 Control Question** **21**

7.1 LPGG . . . 21

7.2 TPGG . . . 21

### 1 Nash equilibria under free allocation

We characterized the pure-strategy Nash equilibria of the one-shot games to provide a theoretical benchmark for the behavioral analyses. Three participants interacted through three dyadic public-goods pools. Each participant received an endowment of $e = 35$ wealth units and allocated contributions, in increments of five units, between the two pools to which they belonged, retaining any unallocated wealth. Let $c^{ij}$ denote participant $i$'s contribution to the pool shared with participant $j$. The feasible strategy set for participant $i$ was

$$\mathcal{S}_i = \left\{(c^{ij}, c^{ik}) : c^{ij}, c^{ik} \in \{0, 5, \ldots, 35\},\ c^{ij} + c^{ik} \leq e\right\}, \tag{1}$$

where $j$ and $k$ denote the other two participants. A feasible contribution profile constituted a Nash equilibrium if no participant could increase their monetary payoff by unilaterally changing the contribution to either or both pools. The analysis assumed that participants maximized their own monetary payoffs and, for the TPGG, that the threshold was fixed and commonly known. We initially abstracted from threshold jitter to distinguish the strategic properties of the game from uncertainty concerning successful provision.

#### 1.1 Linear public goods game

In the LPGG, total contributions to each dyadic pool were multiplied by $r = 1.5$ and divided equally between its two members. Holding all other contributions fixed, an additional unit contributed by the focal participant generated a private return of $r/2 = 0.75$ and therefore reduced their monetary payoff by $1 - r/2 = 0.25$. Zero contribution to both pools was consequently the unique best response to every feasible profile of the other participants' contributions. The LPGG thus had a unique Nash equilibrium:

$$c^{ij} = 0 \qquad \text{for all } i \neq j. \tag{2}$$

Allowing participants to allocate their endowment freely between two pools did not alter this result because the private marginal return remained below the marginal contribution cost in each pool.

#### 1.2 Threshold public goods game

In the TPGG, a pool generates a return only if its multiplied contributions meet or exceed its threshold. For the nominal thresholds $T \in \{30, 45, 60\}$, the required combined contributions are $q = T/r \in \{20, 30, 40\}$, respectively. These requirements are exactly attainable on the experimental contribution grid.

The all-zero profile remains a Nash equilibrium at every nominal threshold. If both partners contribute zero, independently activating a pool requires a contribution of $q$, yielding a net return of $T/2 - q = -T/6 < 0$ relative to retaining that contribution. When $T = 60$, unilateral provision is additionally infeasible because $q = 40 > E$. Thus, no participant benefits from departing unilaterally from the all-zero profile. Unlike in the LPGG, however, zero contribution is not a dominant strategy: contributing can be a best response when a partner supplies enough to make joint provision privately worthwhile.

For the nominal thresholds, a feasible contribution profile is a pure-strategy Nash equilibrium if and only if every dyadic pool satisfies one of two conditions: it is inactive, with both contributions equal to zero; or contributions exactly meeting the provision requirement and neither member contributing more than their return from that pool. Formally, for each unordered pair $\{i, j\}$,

$$
\begin{aligned}
&c_{ij} = c_{ji} = 0, \\
&\text{or} \\
&c_{ij} + c_{ji} = \frac{T}{r}, \qquad 0 < c_{ij} < \frac{T}{2}, \qquad 0 < c_{ji} < \frac{T}{2},
\end{aligned}
\tag{3}
$$

subject to the experimental contribution grid and each participant's joint budget constraint, $\sum_{j \neq i} c_{ij} \leq e$.

Under a fixed and commonly known threshold, replacing the strict inequalities in Eq. 3 with weak inequalities yields the complete set of pure-strategy Nash equilibria. A participant contributing exactly $T/2$ is indifferent between maintaining and withdrawing their contribution because the return from successful provision exactly offsets its cost. Such boundary profiles therefore constitute weak Nash equilibria in the deterministic game. In the experiment, however, threshold jitter introduced a positive probability that a nominally sufficient contribution would fail to generate a return. Under this uncertainty, contributing $T/2$ produced a lower expected payoff than retaining the same amount. We therefore excluded these knife-edge profiles and focused on equilibria that were robust to provision risk, requiring each contributor's cost to be strictly lower than their return from the pool.

These conditions are also sufficient under this refinement. In a threshold-reached pool, reducing any positive contribution causes the total contribution to fall below the threshold, and the foregone return exceeds the contribution saved because $c_{ij} < T/2$. Increasing a contribution raises its cost without increasing its return. In a threshold-not-reached pool, the partner contributes zero, such that unilateral contribution yields a negative net return. Because payoffs are additive across pools, no simultaneous reallocation between the two pools can increase the participant's total payoff when the original profile satisfies the joint budget constraint.

The admissible risk-robust contribution splits for each nominal threshold are shown below. Splits are reported without distinguishing the identities of the two pool members.

| Threshold $T$ | Required total $q$ | Return per member | Admissible robust allocation |
|---|---|---|---|
| 30 | 20 | 15 | $(10, 10)$ |
| 45 | 30 | 22.5 | $(10, 20)$, $(15, 15)$ |
| 60 | 40 | 30 | $(15, 25)$, $(20, 20)$ |

Free allocation couples dyadic contribution decisions through each participant's endowment. At the low and middle thresholds, all three pools can reach the threshold when each participant contributes 10 or 15 units, respectively, to each partner. At the high threshold, provision in all three pools is infeasible because it would require 120 units, exceeding the aggregate endowment of 105 units; nevertheless, equilibria with one or two threshold-reached pools remain possible. The TPGG therefore admits multiple, potentially asymmetric provision patterns, although their existence does not ensure behavioral convergence. Thus, whereas the LPGG makes zero contribution uniquely optimal, the TPGG permits multiple allocation patterns whose feasibility depends on how participants allocate their endowments across partners.

#### 1.3 Supplementary experimental details

Participants completed pre-experiment questionnaires assessing demographic characteristics, economic preferences and social value orientation (Supplementary Section. 5), followed by instructions for their assigned game and comprehension questions (Supplementary Section. 7). The task began only after all comprehension questions had been answered correctly. Following a resting-state recording, participants completed either the linear public goods game (LPGG) or the threshold public goods game (TPGG) in three-person groups, with simultaneous functional near-infrared spectroscopy (fNIRS) recording. Each participant received an endowment of 35 wealth units per round and could allocate it across two dyadic public pools while retaining any unallocated wealth. The experiment comprised six adaptation rounds and 15 main-task rounds. The experimental sequence is illustrated in Supplementary Fig. 1. The behavioral sample comprised 126 participants in 42 groups (21 groups in LPGG and 21 groups in TPGG); the neural sample comprised 120 participants in 40 groups (20 groups in LPGG and 20 groups in TPGG) after exclusions for recording issues.

In the TPGG, a dyadic pool generated a return when its combined contribution, multiplied by 1.5, met or exceeded the realized threshold. The nominal thresholds were 30, 45 and 60 wealth units. Each threshold occurred twice during the adaptation stage and five times during the main task, with presentation order randomized across rounds (Supplementary Fig. 4a). To introduce provision uncertainty, the realized threshold remained at its nominal value with a probability of 0.50, increased by four units with a probability of 0.25 or decreased by four units with a probability of 0.25.

The post-experiment questionnaire assessed participants' preferred contributions, self-reported contribution strategies and perceived accuracy in estimating others' contributions (Supplementary Section. 6). Perceived accuracy was rated from 0 to 6. Reported strategies were classified as conditional, compensatory, prosocial, free-riding or other. Conditional strategies involved adjusting contributions in the same direction as others' contributions, whereas compensatory strategies involved adjustment in the opposite direction. Prosocial and free-riding strategies indicated consistently high and low contributions, respectively.

Free-text explanations of contributing despite possible losses were analysed separately for the LPGG and TPGG using GPT-assisted coding followed by human review. Coding allowed multiple motivational themes within a response. LPGG responses described trust formation, reciprocal contribution and anticipated future benefits, whereas TPGG responses also emphasized threshold feasibility, uncertainty and testing partners' willingness to contribute. These themes describe participants' stated motives and do not independently establish the mechanisms underlying their choices.

### 2 Supplementary behavioral analyses

Post-experiment responses provided complementary evidence on participants' contribution preferences and self-reported strategies (Supplementary Fig. 2). Participants in the LPGG generally reported higher preferred contributions to a dyadic public pool than those in the TPGG, whereas self-rated accuracy in estimating others' contributions was broadly comparable between the two game structures. Conditional contribution was the most frequently reported strategy in both conditions, accounting for 48 of 62 classified responses (77.42%) in the LPGG and 46 of 63 responses (73.02%) in the TPGG. Compensatory, prosocial and free-riding strategies were reported less frequently. Free-riding was more commonly reported in the TPGG (6 of 63 responses, 9.52%) than in the LPGG (3 of 62 responses, 4.84%).

Total individual contributions displayed distinct temporal patterns across the two game structures (Supplementary Fig. 3a). Contributions in the LPGG increased progressively over repeated interaction, rising from approximately 11 wealth units in the initial round to more than 20 units during the later rounds. By contrast, contributions in the TPGG remained comparatively stable and fluctuated around a lower level after the initial rounds. Thus, the divergence between the two game structures emerged gradually, with the LPGG supporting a sustained increase in overall contribution while the TPGG showed no comparable upward trajectory.

Threshold attainment in the TPGG varied markedly across threshold levels (Supplementary Fig. 3b). Across the 21 groups, the mean proportion of dyadic public pools reaching the threshold was highest at the low threshold ($M = 0.460$, SD = 0.241), followed by the middle threshold ($M = 0.254$, SD = 0.202) and the high threshold ($M = 0.077$, SD = 0.110). Attainment was significantly higher at the low than at the middle threshold (paired $t$-test, $t(20) = 5.90$, mean difference = 0.206, 95% CI [0.133, 0.279], $P = 9.05 \times 10^{-6}$), whereas attainment at the high threshold was significantly lower than at the middle threshold ($t(20) = -4.34$, mean difference = $-0.177$, 95% CI [$-0.262$, $-0.092$], $P = 3.21 \times 10^{-4}$). The distributions became increasingly concentrated near zero as the threshold increased, indicating that high-threshold pools were rarely successfully provided under free allocation.

We further examined how contributions evolved across successive encounters with each threshold level (Supplementary Fig. 4b). Contributions increased descriptively across occurrences of the low and middle thresholds, with the clearest increase observed at the middle threshold. By contrast, contributions declined across the early occurrences of the high threshold and remained comparatively low thereafter. This divergence is consistent with adaptation to threshold attainability. At the low and middle thresholds, participants could learn that coordinated contributions had a reasonable probability of generating a return and progressively increase their investment. At the high threshold, however, successful provision required substantially greater joint contributions and was rarely achieved. Repeated failures may therefore have reduced participants' expectations that the pool would reach the threshold, making further contributions increasingly unattractive. These trajectories suggest that thresholds did not provide a uniform coordination signal under free allocation: attainable thresholds could support gradual contribution adjustment, whereas an excessively demanding threshold could instead discourage continued investment.

To assess the robustness of the conditional contribution results, we fitted an alternative Bayesian mixed-effects specification in which random intercepts and random slopes for the focal participant's and partner's preceding contributions were defined at the dyadic-public-pool level. This differs from the primary analysis reported in the main text, in which the corresponding random effects were grouped by individual identity. The pool-level model reproduced the main pattern: both stickiness and imitation were positive in each game and stronger in the LPGG (stickiness: posterior mean $\beta = 0.38$, SE = 0.02, 95% credible interval [0.33, 0.43]; imitation: $\beta = 0.45$, SE = 0.03, 95%

credible interval [0.40, 0.51]) than in the TPGG (stickiness: $\beta$ = 0.28, SE = 0.03, 95% credible interval [0.22, 0.34]; imitation: $\beta$ = 0.27, SE = 0.03, 95% credible interval [0.22, 0.33]; Supplementary Fig. 5a,b).

We additionally examined whether round-to-round changes in the focal participant's contribution covaried with contemporaneous changes in the partner's contribution to the same dyadic public pool (Supplementary Fig. 5c,d). Bayesian mixed-effects models included participant-specific random intercepts and random slopes for the partner's contribution change. Partner contribution changes positively predicted focal contribution changes in both games, with a larger posterior estimate in the LPGG ($\beta$ = 0.36, SE = 0.04, 95% credible interval [0.28, 0.43]) than in the TPGG ($\beta$ = 0.14, SE = 0.04, 95% credible interval [0.06, 0.22]). This difference may arise because the continuous payoff structure of the LPGG preserves the behavioral relevance of incremental changes in a partner's contribution, allowing participants to respond proportionally to partner-specific adjustments. In the TPGG, by contrast, the consequences of a contribution change depend on the current threshold and the partner's contribution, such that an equivalent adjustment may either leave provision unchanged or determine whether the threshold is reached. This nonlinear mapping may reduce the informational value of incremental contribution changes and thereby weaken direct behavioral matching between partners.

### 3 Supplementary neural analyses

#### 3.1 fNIRS optode configuration and cortical coverage

Neural signals were recorded simultaneously from all three group members using identical optode configurations positioned according to the international 10–10 system (Supplementary Fig. 6). The frontal array comprised five emitters and four detectors, forming 12 measurement channels over the dorsomedial prefrontal cortex (dmPFC). An additional emitter–detector pair positioned at CP6 and TP8 formed one channel over the right temporoparietal junction (rTPJ). Optode positions and signal quality were verified before each experimental session. Subsequent analyses were based on changes in oxygenated haemoglobin concentration.

#### 3.2 Selection of the frequency band for inter-brain neural synchrony

We first identified the frequency range in which inter-brain neural synchrony was most strongly modulated during task performance. Wavelet transform coherence was averaged across participant pairs and the 13 channels, and task-related changes were calculated relative to the pre-task resting-state baseline (Supplementary Fig. 7). This analysis revealed an increase in coherence within the 0.0288–0.0384-Hz range, corresponding to oscillation periods of approximately 26–35 s. This interval closely matched the duration of the task-related sequence, comprising the 12-s operation phase and the subsequent 13–23-s outcome and waiting period.

The identified frequency range lies below the bands typically dominated by respiratory, cardiac and rapid motion-related fluctuations. We therefore defined 0.0288–0.0384 Hz as the frequency band of interest and restricted all subsequent INS analyses to coherence values within this interval. For each dyad and channel, task-related INS was quantified as the Fisher-z-transformed coherence during the operation phase relative to the resting-state baseline.

#### 3.3 Additional analysis of association between INS and behavior

To examine how the association between dyadic contribution differences and inter-brain neural synchrony (INS) developed over repeated interaction, we conducted a cumulative-window analysis in the LPGG (Supplementary Fig. 8). Beginning with rounds 1–7, we progressively extended the analysis window by one round until all 21 rounds were included. Within each window, contribution differences and Channel-3 dmPFC INS were averaged across the included rounds for each dyad, and their association was estimated using linear regression.

The estimated association was consistently positive across all cumulative windows ($\beta > 0$), indicating that dyads with greater contribution differences exhibited stronger Channel-3 INS. The effect increased during the early and middle stages of the interaction, reached its largest magnitude in windows ending around rounds 13–15 and subsequently declined modestly while remaining positive. In the full 21-round window, the association remained significant ($\beta = 0.016$, SE $= 0.008$, $t(58) = 2.10$, $F(1, 58) = 4.40$, $P = 0.0404$). This temporal pattern suggests that dmPFC synchrony tracked the cumulative development of partner-specific contribution differences rather than a transient effect confined to a particular phase of the task.

Because successive cumulative windows contained overlapping observations, the estimates should not be interpreted as independent replications. This analysis was therefore used to characterize the temporal emergence of the association rather than to provide a series of independent confirmatory tests.

### 4 Analysis of post-experiment measures

#### 4.1 GPT-assisted qualitative analysis of post-experiment free-text responses

We used a GPT-assisted qualitative text-analysis procedure to characterize participants' stated reasons for contributing despite the possibility of incurring losses. Free-text responses were analysed separately for the linear public goods game (LPGG; n = 63) and the threshold public goods game (TPGG; n = 63). Responses were semantically coded into non-mutually exclusive motivational themes, allowing each response to reflect multiple motives. The resulting codes were then reviewed and consolidated into interpretable categories for each game type.

In the TPGG, participants' explanations were more explicitly conditional on whether the collective threshold could be reached. Contributions were commonly described as attempts to coordinate with others, test partners' willingness to cooperate, signal commitment, and form stable cooperative relationships that would enable the group to exceed the threshold and obtain higher payoffs. Compared with the LPGG condition, TPGG responses placed greater emphasis on threshold feasibility, risk assessment, partner screening and strategic trial-and-error. Several participants also indicated that they would contribute only when the threshold was sufficiently low, when others appeared likely to contribute, or when the expected return outweighed the potential loss. Thus, in the threshold public goods setting, costly contribution was primarily motivated by threshold-oriented coordination, conditional cooperation and strategic exploration of others' cooperativeness.

Across both game types, participants rarely described costly contribution as purely unconditional altruism. Instead, contributions were generally understood as strategic investments aimed at eliciting reciprocity, establishing trust, coordinating future behavior and increasing long-term payoffs. However, the motivational emphasis differed between the two institutional structures: LPGG responses centred on trust formation and reciprocal cooperation under continuous returns, whereas TPGG responses centred on reaching a collective threshold under uncertainty, with stronger attention to risk, feasibility and partner selection.

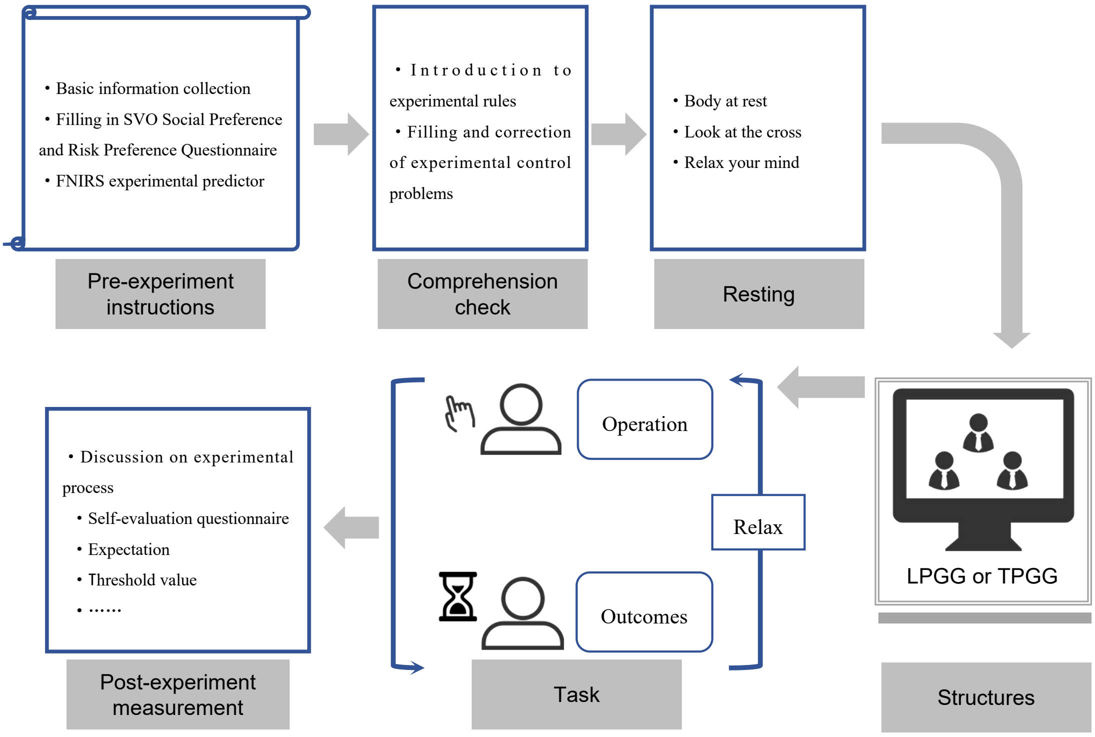

**Supplementary Fig. 1: Overview of the experimental procedure**. Participants completed pre-experiment questionnaires assessing demographic characteristics, social value orientation and economic preferences, followed by instructions and comprehension checks for their assigned game. All comprehension questions had to be answered correctly before the task began. After a resting-state recording, participants played either the LPGG or TPGG in three-person groups while functional near-infrared spectroscopy (fNIRS) signals were recorded simultaneously. Each participant could allocate their endowment across two dyadic public pools and retain any unallocated wealth. The task comprised 21 rounds, including six adaptation rounds and 15 main-task rounds, with repeated decision, outcome and rest periods. Post-experiment questionnaires assessed contribution preferences, self-reported strategies and beliefs about other participants' behavior.

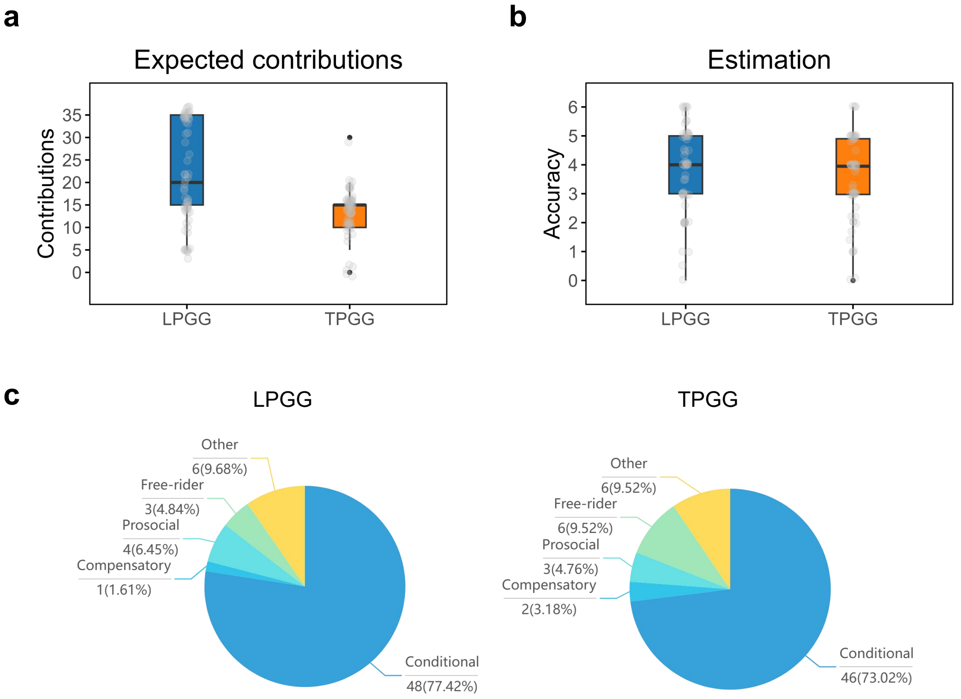

**Supplementary Fig. 2**: **Self-reported contribution expectations, estimation accuracy and behavioral strategies. a,** Expected contribution to a dyadic public pool reported after completion of the LPGG or TPGG. **b,** Self-rated accuracy in estimating other participants' contributions, assessed on a scale from 0 (completely inaccurate) to 6 (completely accurate). In **a** and **b**, grey points represent individual responses ($n = 63$ participants per game structure). **c,** Distribution of self-reported contribution strategies in the LPGG ($n = 62$ classified responses) and TPGG ($n = 63$ classified responses). Conditional strategies involved increasing contributions when others contributed more and decreasing them when others contributed less, whereas compensatory strategies followed the opposite pattern. Prosocial and free-riding strategies reflected consistently high and low contributions, respectively, irrespective of others' behavior. Labels indicate the number and percentage of responses assigned to each category.

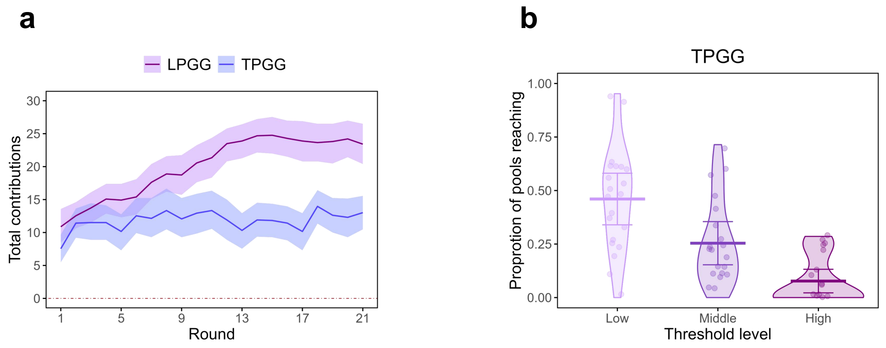

**Supplementary Fig. 3**: **Contribution dynamics and threshold attainment. a,** Mean total individual contributions across 21 rounds in the LPGG (purple) and TPGG (blue). Total contribution was defined as the sum allocated by an individual to their two dyadic public pools in each round. Shaded bands represent the s.e.m. across participants ($n = 63$ per game structure). **b,** Distribution of the proportion of dyadic public pools reaching the threshold at each threshold level in the TPGG. Points represent individual three-person groups ($n = 21$), violin plots show the corresponding distributions, and horizontal lines and error bars indicate the mean and 95% confidence interval, respectively.

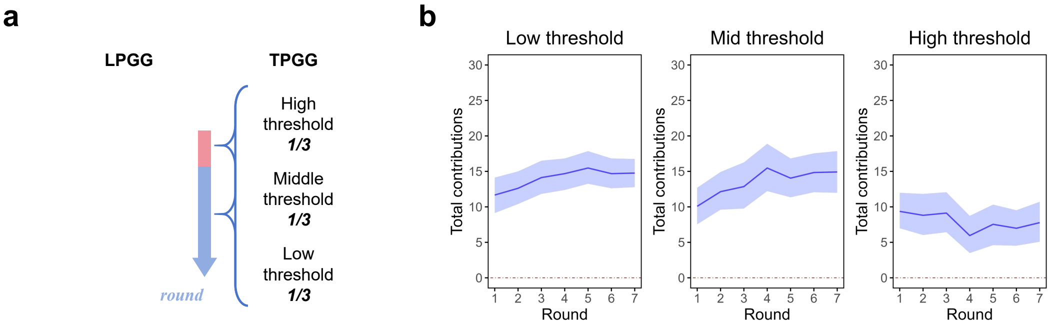

**Supplementary Fig. 4**: **Contribution trajectories across threshold levels. a,** Schematic of threshold presentation in the TPGG. The low, middle and high nominal thresholds were 30, 45 and 60 wealth units, respectively, with each level occurring in one-third of the 21 rounds. Each threshold level appeared twice during the adaptation stage and five times during the main task, in randomized order. The LPGG had no provision threshold. **b,** Mean total individual contributions across the seven occurrences of each threshold level in the TPGG. The horizontal axis denotes the successive occurrence of a given threshold level rather than consecutive rounds of the full experiment. Solid lines indicate means, and shaded bands represent the s.e.m. across participants ($n = 63$).

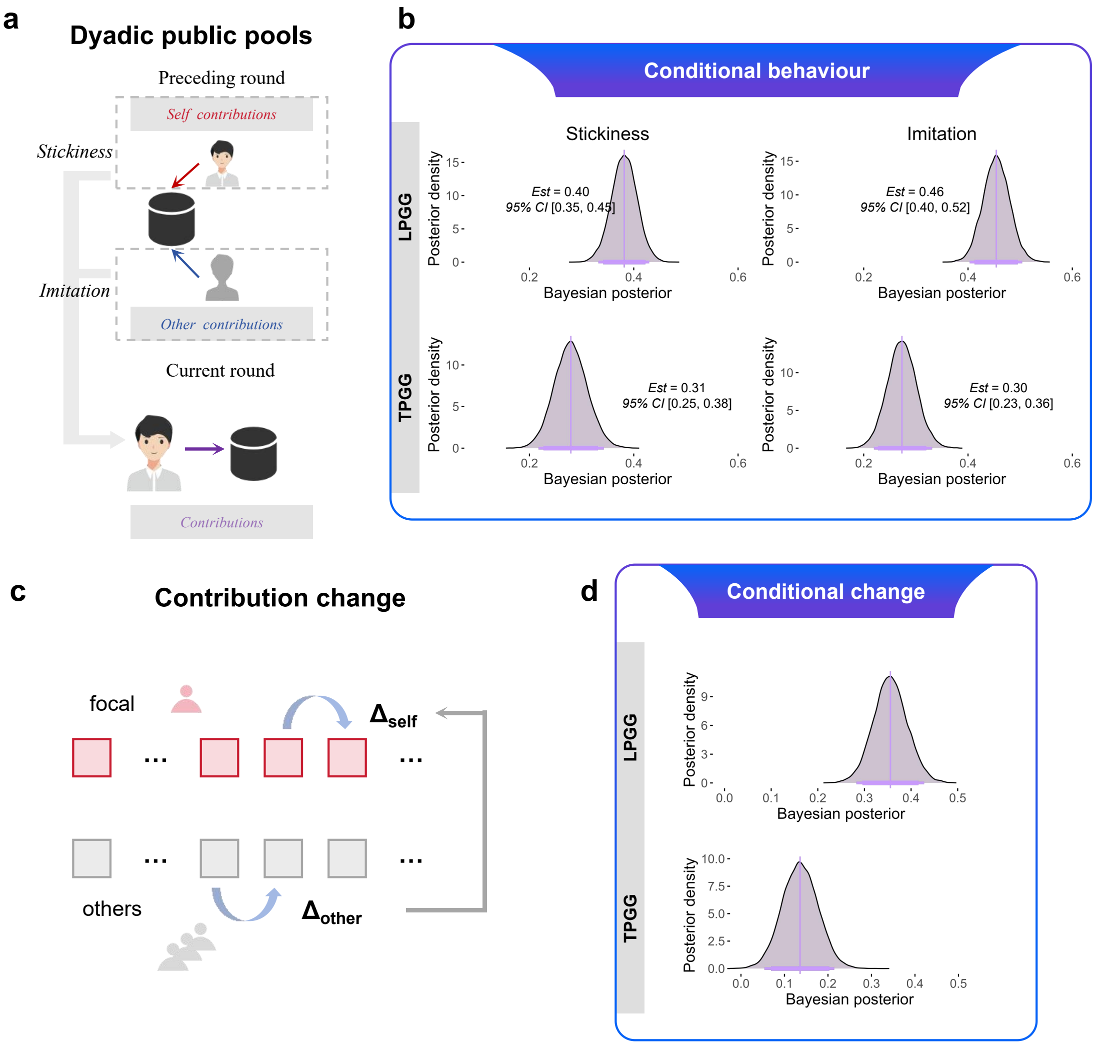

**Supplementary Fig. 5: Conditional contributions within dyadic public pools and conditional contribution change. a,** Schematic of the conditional contribution analysis. The focal participant's contribution to a dyadic public pool in the current round was modelled as a function of their own and their partner's contributions to the same pool in the preceding round. The corresponding coefficients quantify stickiness and imitation, respectively. **b,** Posterior distributions of the stickiness and imitation coefficients estimated separately for the LPGG and TPGG. **c,** Schematic of the conditional contribution change analysis, relating the focal participant's round-to-round change in contribution to the corresponding change in their partner's contribution within the same dyadic pool. **d,** Posterior distributions of the coefficients linking partner and focal contribution changes in each game structure. Models were fitted separately for each game and included individual-specific random intercepts and random slopes.

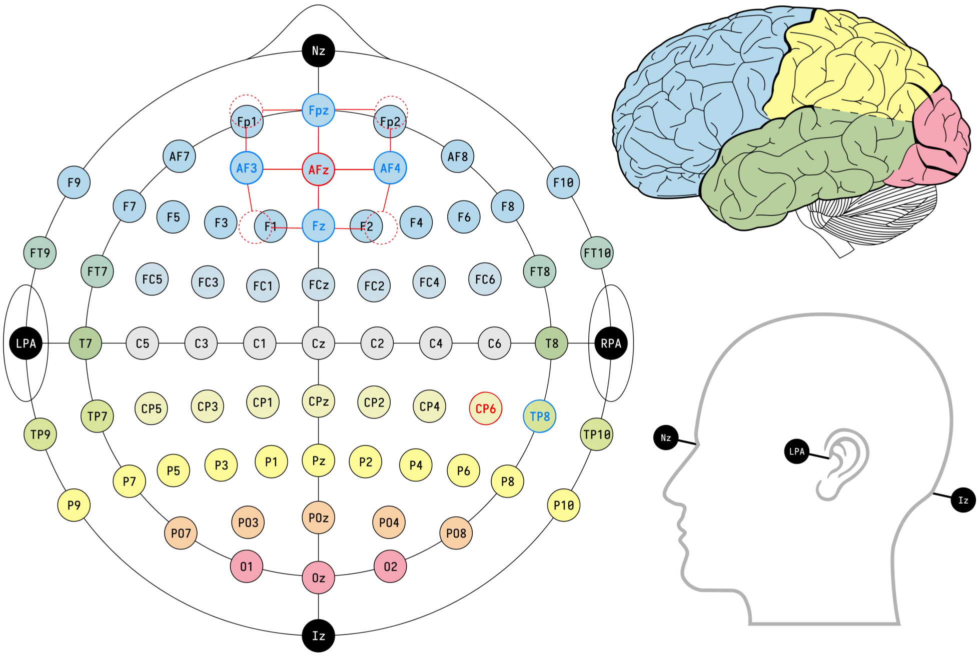

**Supplementary Fig. 6**: **fNIRS optode placement and anatomical reference landmarks.** Optode positions are mapped onto the international 10–10 scalp coordinate system. The frontal array covered the dorsomedial prefrontal cortex and comprised five emitters and four detectors, forming 12 measurement channels. The right temporoparietal array comprised one emitter positioned at CP6 and one detector positioned at TP8, forming one additional channel. Red and blue labels denote emitter and detector positions, respectively, and connecting lines indicate measurement channels. The same optode arrangement was used for all three participants in each group.

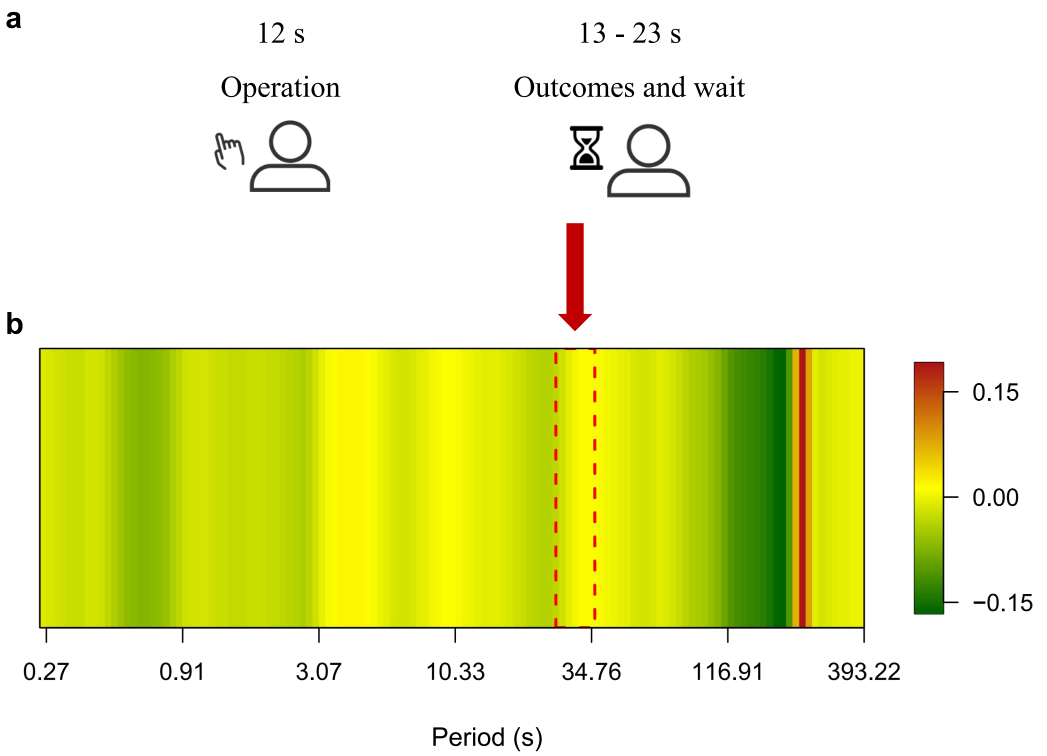

**Supplementary Fig. 7**: **Task timing and frequency-band selection for inter-brain neural synchrony. a,** Schematic of the task-related time window, comprising a 12-s operation phase followed by a 13–23-s outcome and waiting period. **b,** Task-related changes in Fisher-z-transformed wavelet coherence relative to the pre-task resting-state baseline, averaged across participant pairs and the 13 channels and plotted as a function of oscillation period. Colours indicate the magnitude and direction of the change in coherence. Red dashed lines delimit the frequency band selected for subsequent inter-brain neural synchrony analyses (0.0288–0.0384 Hz), corresponding to periods of approximately 26–35 s; the red arrow highlights this interval.

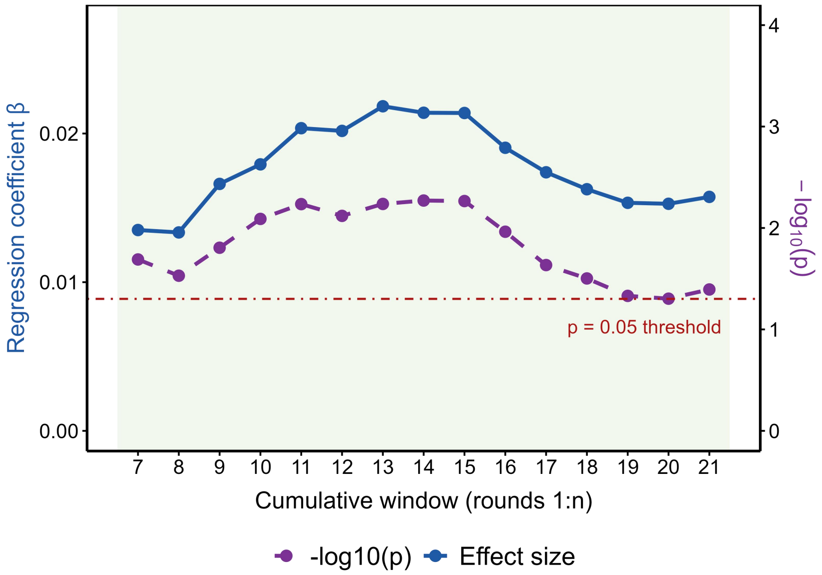

**Supplementary Fig. 8**: **Cumulative-window analysis of the association between contribution differences and inter-brain neural synchrony.** The association between mean dyadic contribution differences and Channel-3 inter-brain neural synchrony in the LPGG was estimated over cumulative windows extending from round 1 to round $n$, where $n$ ranged from 7 to 21. For each window, behavioral and neural measures were averaged across the included rounds before fitting a linear regression model. The solid blue line shows the estimated regression coefficient, $\beta$ (left axis), and the dashed purple line shows $-\log_{10}(P)$ (right axis). The horizontal red dash-dotted line denotes the nominal significance threshold of $P = 0.05$.

#### 5 Pre-experiment questionnaire

1. Name [Fill-in-the-blank question] *

________________

2. Date of Birth [Fill-in-the-blank question] *

________________

3. Gender [Single-choice question] *

[ ]Male [ ]Female

The following statements describe what kind of person you are. Please indicate a number from 0 to 10——0 means [This statement does not describe me at all], and 10 means [This statement perfectly describes me]. You can also use any number between 0 and 10 to represent your position on this scale, such as 0, 1, 2, 3, 4, 5, 6, 7, 8, 9, 10.

4. Compared with others, I am a person who is willing to punish unfair behavior even if the cost of punishment is high. [Enter a number from 0 (does not match) to 10 (matches)] *

________________

5. I make a special effort to help those who have helped me before. [Enter a number from 0 (does not match) to 10 (matches)] *

________________

6. If someone is unfair to me in sports, I will be unfair to them in return. [Enter a number from 0 (does not match) to 10 (matches)] *

________________

7. I am willing to spend time and money on non - profit things that seem meaningful to me, even if they do not directly benefit me. [Enter a number from 0 (does not match) to 10 (matches)] *

________________

8. If someone does me a favor, I am willing to repay them. [Enter a number from 0 (does not match) to 10 (matches)] *

________________

9. If I suffer a serious injustice, I will take revenge at all costs next time I have the chance. [Enter a number from 0 (does not match) to 10 (matches)] *

________________

10. I am willing to help others, even if I think I will never meet them again. [Enter a number from 0 (does not match) to 10 (matches)] *

________________

11. I don't understand why some people spend their whole lives fighting for a cause that doesn't directly benefit them. [Enter a number from 0 (does not match) to 10 (matches)] *

________________

12. Compared with others, I am a person who is willing to take risks. [Enter a number from 0 (does not match) to 10 (matches)] *

________________

13. When others are in trouble or an emergency, I am willing to face the difficulties with them without expecting any return. [Enter a number from 0 (does not match) to 10 (matches)] *

________________

14. Compared with others, I am a person who usually likes to share with others without expecting something in return. [Enter a number from 0 (does not match) to 10 (matches)] *

________________

The following are hypothetical scenario questions.

15. Imagine: I got lost while shopping in a strange city and asked a stranger for directions. The stranger offered to drive me to my destination. The drive took about 20 minutes, and the stranger spent about 100 yuan in total. The stranger didn't want money. I had six bottles of wine with me, with the following prices. I decided to give the stranger a bottle of wine as a thank - you gift. [Single-choice question] *

[ ]25 [ ]50 [ ]75 [ ]100 [ ]125 [ ]150

16. Imagine: I won 10,000 yuan in the lottery. Considering my current situation, I would donate ___________ yuan to a charity. [Fill-in-the-blank question] *

17. Imagine: I participated in an activity with a stranger and we won 100 yuan. The rule is that the stranger proposes a plan to divide the 100 yuan. After I know the plan, I decide whether to accept or reject it. If I accept, the money will be divided according to the plan; if I reject, neither of us will get anything. After thinking, I think the other party should offer me at least ___________ yuan for me to accept. [Fill-in-the-blank question] *

18. Imagine: I won a competition and can choose between two payment methods: (1) a lottery where I have a 50% chance of getting 1000 yuan and a 50% chance of getting nothing; (2) a fixed bonus between 0 and 1000 yuan. The fixed bonus should be at least ___________ yuan for me to prefer it to the lottery. [Fill-in-the-blank question] *

Imagine: I am doing an activity with a stranger. The rule is that we each start with 20 yuan. First, the other party can transfer any integer amount from 0 to 20 yuan to me, that is, 0, 1, 2, ..., up to 20. The amount transferred to me will be tripled and added to my account. Then I have the opportunity to transfer any integer amount back to the other party. The amount does not double in this process, and the maximum amount I can transfer depends on the total amount in my account after receiving the transfer from the other party.

19. Suppose the other party transfers 5 yuan to me. I have 20 + 3×5 = 35 yuan, and the other party has 20-5=15 yuan. I decide to transfer back ___________ yuan. [Fill-in-the-blank question] *

20. Suppose the other party transfers 10 yuan to me. I have 20 + 3×10 = 50 yuan, and the other party has 20 - 10 = 10 yuan. I decide to transfer back ___________ yuan. [Fill-in-the-blank question] *

21. Suppose the other party transfers 15 yuan to me. I have 20 + 3×15 = 65 yuan, and the other party has 20 - 15 = 5 yuan. I decide to transfer back ___________ yuan. [Fill-in-the-blank question] *

22. Suppose the other party transfers 20 yuan to me. I have 20 + 3×20 = 80 yuan, and the other party has 20 - 20 = 0 yuan. I decide to transfer back ___________ yuan. [Fill-in-the-blank question] *

23. Suppose I were in the other person's party. First, I will transfer ___________ yuan. [Fill-in-the-blank question] *

Suppose you receive the following series of plans. For each plan, you need to select your favorite way of distribution to divide the wealth between you and another stranger.

### 6 Post-experiment measure

Dear students, please fill out the questionnaire carefully. If you have any questions, feel free to ask the experimenter in a timely manner.

1. How much personal wealth do you tend to contribute to a dyadic community? [Single choice] *

[ ]0 [ ]5 [ ]10 [ ]15 [ ]20 [ ]25 [ ]30 [ ]35

2. Your contribution strategy (rank multiple options by importance; enter numbers in brackets accordingly): [Ranking] *

[ ] If other participants contribute more, I contribute more; if they contribute less, I contribute less [ ] If other participants contribute less, I contribute more; if they contribute more, I contribute less [ ] Always maintain a low contribution regardless of others [ ] Always maintain a high contribution regardless of others [ ] Contribute randomly [ ] Other

3. Given that contributions may result in a loss, what is your motivation for contributing? [Open-ended] *

____________________

4. During the experiment, did you attempt to estimate other participants' contributions? [Single choice] *

[ ]Yes [ ]No

5. How accurate do you think your estimates were? [Enter a number from 0 (inaccurate) to 6 (precise)] *

____________________

6. (Specific to TPGG) What methods did you use to reach the threshold during the experiment? [Open-ended] *

____________________

### 7 Control Question

#### 7.1 LPGG

To ensure that participants understood the experimental rules, they answered the following control questions by entering their calculated results in the input box, in wealth points.

The initial endowment in each round was 35. Suppose that the productivity factor was 1.2. Note that this differs from the productivity factor of 1.5 used in the experiment.

(1) Suppose that all participants contribute 0 wealth points.

Question 1: What is your final wealth?

Question 2: What is the sum of the final wealth of the other two participants?

(2) Suppose that participant 2 contributes 0 wealth points to community 1–2 and participant 3 contributes 0 wealth points to community 1–3.

Question 3: In addition, you contribute 10 wealth points only to community 1–2. What is your final wealth?

Question 4: In addition, you contribute 15 wealth points only to community 1–2. What is your final wealth?

Question 5: In addition, you contribute 20 wealth points only to community 1–2. What is your final wealth?

Question 6: In addition, you contribute 15 wealth points to each of communities 1–2 and 1–3. What is your final wealth?

(3) Suppose that participant 2 contributes 20 wealth points to community 1–2 and participant 3 contributes 20 wealth points to community 1–3.

Question 7: In addition, you make no contribution. What is your final wealth?

Question 8: In addition, you contribute 15 wealth points only to community 1–2. What is your final wealth?

Question 9: In addition, you contribute 15 wealth points only to community 1–3. What is your final wealth?

Question 10: In addition, you contribute 20 wealth points only to community 1–3. What is your final wealth?

Question 11: In addition, you contribute 15 wealth points to each of communities 1–2 and 1–3. What is your final wealth?

(4) Suppose that you contribute 5 and 10 wealth points to communities 1–2 and 1–3, respectively.

Question 12: In addition, participant 2 contributes 20 wealth points to community 1–2 and participant 3 contributes 5 wealth points to community 1–3. What is your final wealth?

Question 13: In addition, participant 2 contributes 5 wealth points to community 1–2 and participant 3 contributes 20 wealth points to community 1–3. What is your final wealth?

Question 14: In addition, participant 2 contributes 20 wealth points to community 1–2 and participant 3 contributes 15 wealth points to community 1–3. What is your final wealth?

Question 15: In addition, participant 2 contributes 10 wealth points to community 1–2 and participant 3 contributes 15 wealth points to community 1–3. What is your final wealth?

#### 7.2 TPGG

To ensure that participants understood the experimental rules, they answered the following control questions by entering their calculated results in the input box, in wealth points.

The initial endowment in each round was 35. Suppose that the productivity factor was 1.2. Note that this differs from the productivity factor of 1.5 used in the experiment.

(1) Suppose that all participants contribute 0 wealth points.

Question 1: What is your final wealth?

Question 2: What is the sum of the final wealth of the other two participants?

(2) Suppose that the thresholds of communities 1–2 and 1–3 are both 48, participant 2 contributes 15 wealth points to community 1–2, and participant 3 contributes 20 wealth points to community 1–3.

Question 3: In addition, you contribute 20 wealth points only to community 1–2. What is your final wealth?

Question 4: In addition, you contribute 20 wealth points only to community 1–3. What is your final wealth?

Question 5: In addition, you contribute 15 wealth points only to community 1–2. What is your final wealth?

Question 6: In addition, you contribute 20 and 15 wealth points to communities 1–2 and 1–3, respectively. What is your final wealth?

(3) Suppose that the thresholds of communities 1–2 and 1–3 are both 36, participant 2 contributes 10 wealth points to community 1–2, and participant 3 contributes 20 wealth points to community 1–3.

Question 7: In addition, you make no contribution. What is your final wealth?

Question 8: In addition, you contribute 15 wealth points only to community 1–2. What is your final wealth?

Question 9: In addition, you contribute 20 wealth points only to community 1–2. What is your final wealth?

Question 10: In addition, you contribute 15 wealth points only to community 1–3. What is your final wealth?

Question 11: In addition, you contribute 15 wealth points to each of communities 1–2 and 1–3. What is your final wealth?

(4) Suppose that the thresholds of communities 1–2 and 1–3 are both 72, and that you contribute 30 and 5 wealth points to communities 1–2 and 1–3, respectively.

Question 12: In addition, participant 2 contributes 30 wealth points only to community 1–2, and participant 3 contributes 25 wealth points only to community 1–3. What is your final wealth?

Question 13: In addition, participant 2 contributes 25 wealth points only to community 1–2, and participant 3 contributes 30 wealth points only to community 1–3. What is your final wealth?

Question 14: In addition, participant 2 contributes 35 wealth points only to community 1–2, and participant 3 contributes 35 wealth points only to community 1–3. What is your final wealth?

### References

[1] Hilbe, C., Chatterjee, K. & Nowak, M. A. Partners and rivals in direct reciprocity. *Nature human behaviour* **2**, 469–477 (2018).

[2] Wang, X. *et al.* Cooperation and coordination in heterogeneous populations. *Philosophical Transactions of the Royal Society B: Biological Sciences* **378** (2023).

[3] Park, S. A., Sestito, M., Boorman, E. D. & Dreher, J.-C. Neural computations underlying strategic social decision-making in groups. *Nature communications* **10**, 5287 (2019).
